# Distinct regionalization patterns of cortical morphology are associated with cognitive performance across different domains

**DOI:** 10.1101/2020.02.13.948596

**Authors:** C E Palmer, W Zhao, R Loughnan, J Zou, C C Fan, W K Thompson, A M Dale, T L Jernigan

**Affiliations:** Center for Human Development, University of California, San Diego, 9500 Gilman Drive, La Jolla, CA 92161, USA; Department of Cognitive Science, University of California, San Diego, 9500 Gilman Drive, La Jolla, CA 92093, USA; Center for Multimodal Imaging and Genetics, University of California, San Diego School of Medicine, 9444 Medical Center Dr, La Jolla, CA 92037, USA; Division of Biostatistics, Department of Family Medicine and Public Health, University of California, San Diego, La Jolla, CA, USA; Department of Radiology, University of California, San Diego School of Medicine, 9500 Gilman Drive, La Jolla, CA 92037, USA; Department of Neuroscience, University of California, San Diego School of Medicine, 9500 Gilman Drive, La Jolla, CA 92037, USA; Department of Psychiatry, University of California, San Diego School of Medicine, 9500 Gilman Drive, La Jolla, CA 92037, USA

**Author notes:** Corresponding author: Dr. Clare E Palmer.

**Keywords:** adolescence, cognition, cortical morphology, development, multivariate, neuroimaging

## Abstract

Cognitive performance in children is predictive of academic and social outcomes; therefore, understanding neurobiological mechanisms underlying individual differences in cognition during development may be important for improving quality of life. The belief that a single, psychological construct underlies many cognitive processes is pervasive throughout society. However, it is unclear if there is a consistent neural substrate underlying many cognitive processes. Here we show that a distributed configuration of cortical surface area and apparent thickness, when controlling for global imaging measures, is differentially associated with cognitive performance on different types of tasks in a large sample (N=10,145) of 9-11 year old children from the Adolescent Brain and Cognitive Development^SM^ (ABCD) study. The minimal overlap in these regionalization patterns of association has implications for competing theories about developing intellectual functions. Surprisingly, *not* controlling for sociodemographic factors increased the similarity between these regionalization patterns. This highlights the importance of understanding the shared variance between sociodemographic factors, cognition and brain structure, particularly with a population-based sample such as ABCD.

## INTRODUCTION

Two-factor theories of intellectual development have divided cognitive function into two broad components: fluid abilities refer to one’s propensity to solve problems, reason, act quickly and adapt to novel situations; whereas, crystallised abilities encompass task-specific knowledge that accrues throughout the lifespan (Horn and Cattell 1966; Deary 2012). Despite these constructs being seemingly dissociable, performance across different tasks probing these cognitive processes is often correlated (Spearman 1904; Wechsler 1946; Akshoomoff et al. 2013). The latent factor explaining this shared variance, termed ‘g’, (Gottfredson and Deary 2004; Arendt and Nielsen 2005; Cutler et al. 2012) is heritable(Bouchard and McGue 1981; Gottfredson and Deary 2004; Plomin and Spinath 2004; Panizzon et al. 2014) and predictive of social and academic outcomes (Gottfredson and Deary 2004; Arendt and Nielsen 2005; Cutler et al. 2012). A pervasive view in society is that ‘g’ represents a single causal underlying trait that influences performance across multiple domains. Although genome-wide-association-studies (GWAS) demonstrate that cognitive phenotypes, including ‘g’, can be associated with specific genetic variants (Davies et al. 2018; Savage et al. 2018), some of which are also associated with brain structure (Grasby et al. 2020; Lett et al. 2020), these associations are highly polygenic, and though the mediating pathway from genetics to cognition is still poorly understood, there is little support for a causal pathway associated with a unitary neural correlate. Cognitive development is highly complex and some models of intellectual development have posited that correlations in performance across multiple tasks, the so-called “positive manifold”, may emerge via positive interactions among developing cognitive systems, which may have distinct neural substrates (Van Der Maas et al. 2006, 2017; Hampshire et al. 2012). This framework is difficult to test empirically, however one prediction from this mutualism model would be that individual differences in cognitive performance would be associated with different distributions of neural phenotypes across individuals.

Previous studies have reported associations of general cognitive ability with larger brain volume and distributed structural and functional neural correlates particularly across frontal and parietal regions (Duncan et al. 2000; Colom et al. 2009, 2013; Gläscher et al. 2010; Basten et al. 2015). Cortical volume, as measured with MRI, is comprised of two genetically and developmentally distinct components, surface area (CSA) and apparent thickness (CTH), (Panizzon et al. 2009; Winkler et al. 2010; Jernigan et al. 2011), which have shown different associations with individual variability in cognition. Positive associations with cortical surface area (CSA) and cognition have been consistently reported (Schnack et al. 2015; Walhovd et al. 2016; Schmitt et al. 2019; Grasby et al. 2020); however, associations between cortical thickness (CTH) and cognition have been less consistent (Sowell et al. 2004; Shaw et al. 2006; Brouwer et al. 2014; Burgaleta et al. 2014). This is likely due to limited statistical power to identify replicable associations, differences in neuroimaging processing protocols and cognitive assessments used to estimate general cognitive ability, and the age of the participants. Few studies have measured individual differences in the regionalization of cortical morphology (i.e., the relative increase or decrease in CTH or CSA within a particular region relative to mean CTH or total CSA respectively). Complex patterns of positive and negative associations of relative CSA and CTH have been associated with general cognitive function (Fjell et al. 2015; Vuoksimaa et al. 2016; Reardon et al. 2018), as well as with more specific measures of cognitive performance (Fjell et al. 2012; Newman, Jernigan, et al. 2016; Newman, Thompson, et al. 2016; Curley et al. 2018). However, many previous studies have been underpowered to detect significant effects of relative cortical configuration across the whole cortex particularly when using univariate statistical methods with stringent control for multiple comparisons (Vuoksimaa et al. 2016). However, we know that during embryonic development, the relative areal expanse associated with different functional regions occurs via the graded expression of transcription factors across the cortical plate (O’Leary et al. 2007; Rakic et al. 2009). Individual variability in this patterning could therefore result in subtle changes to the whole configuration of the cortex, which may in turn influence cognitive development.

In the current study, we used a multivariate analysis to measure the association between the regionalization of cortical surface area (CSA) and apparent cortical thickness (CTH) in a large sample (n=10,145) of 9-11 year old children from the Adolescent Brain and Cognitive Development^SM^ (ABCD) Study. This large-scale study of 11,880 nine and ten-year-old children, uses neuroimaging, genetics and a multi-dimensional battery of assessments to investigate the role of various biological, environmental, and behavioral factors in brain, cognitive, and social/emotional development. Given the biology of cortical development and growing evidence that models encompassing distributed behavioral associations across the brain better predict behavioral outcomes (Reddan et al. 2017; van der Meer et al. 2020; Zhao et al. 2020), we used a multivariate approach to assess the significance of the effects of the brain phenotypes on behavior when aggregated across all cortical vertices. We subsequently examined the degree to which the distributed cortical patterns associating with cognitive performance measured with the fluid and crystallized composite scores from the NIH Toolbox^®^ exhibited effects of a common underlying cortical architecture or distinct patterns of cortical regionalization. These composite scores generated from the Toolbox have been validated against gold standard measures of intelligence in both adults and children (Akshoomoff et al. 2013; Hodes et al. 2013; Heaton et al. 2014).

The ABCD Study^®^ sample was recruited to resemble the population of the United States as closely as possible; therefore, the participants are from diverse racial, ethnic and socioeconomic backgrounds. Confounding associations between these sociodemographic variables, brain structure and cognitive performance complicate the interpretation of model results that control for demographic covariates. We have therefore performed our analyses with and without controlling for these important sociodemographic variables in order to highlight the implications of this for the interpretation of our findings. This is an important aspect of analyzing individual differences in large population-based samples such as the ABCD Study^®^.

## Materials & Methods

### Sample

The ABCD study is a longitudinal study across 21 data acquisition sites enrolling 11,880 children starting at 9-11 years old. This paper analyzed the baseline sample from release 2.0.1 (NDAR DOI: 10.15154/1504041). The ABCD Study used school-based recruitment strategies to create a population-based, demographically diverse sample, however it is not necessarily representative of the U.S. national population (Garavan et al. 2018; Compton et al. 2019). Due to the inclusion of a wide range of individuals across different races, ethnicities and socioeconomic backgrounds, it is important to carefully consider how to control for these potentially confounding factors and the implications of this on our effects of interest. Sex and age were used as covariates in all analyses. Only subjects who had complete data across all of the measures analyzed were included in the neuroimaging analyses creating a final sample of 10,145 subjects. Additional sample details can be found in the Supplementary Material. Supplementary Table 1 displays the names of each variable used in these analyses from data release 2.0.1. Supplementary Table 2 shows the demographic characteristics of the sample.

### Neurocognitive assessment

The ABCD Study neurocognitive assessment at baseline consisted of the NIH Toolbox Cognition Battery^®^ (NIHTBXCB), the WISC-V matrix reasoning task, the Little Man Task (LMT) and the Rey Auditory Verbal Learning task (RAVLT). All of the tasks were administered using an iPad with support or scoring from a research assistant where needed. The NIHTBXCB is a widely used battery of cognitive tests that measures a range of different cognitive domains. It includes the Toolbox Oral Reading Recognition Task^®^ (TORRT), Toolbox Picture Vocabulary Task^®^ (TPVT), the Toolbox Pattern Comparison Processing Speed Test^®^ (TPCPST), Toolbox List Sorting Working Memory Test^®^ (TLSWMT), Toolbox Picture Sequence Memory Test^®^ (TPSMT), Toolbox Flanker Task^®^ (TFT) and Toolbox Dimensional Change Card Sort Task^®^ (TDCCS). Details of each task can be found in supplementary materials. In the current study, the uncorrected scores for each task were used for statistical analyses. Composite scores estimating crystallized intelligence (mean of TPVT and TORRT), fluid intelligence (mean of TPCPST, TLSWMT, TPSMT, TFT and TDCCS) and a total cognition score (mean of all tasks) are also provided by the NIHTBCB (https://www.healthmeasures.net/explore-measurement-systems/nih-toolbox/intro-to-nih-toolbox/cognition) and were also analyzed. These measures have been validated against ‘gold standard’ measures of general cognitive ability, and the constructs referred to as crystallized and fluid intelligence in adults(Heaton et al. 2014) and children(Akshoomoff et al. 2013). To address remaining concern that the Total Composite Score may not adequately reflect estimates of’g’ as a global latent factor, we conducted a Principal Component Analysis (PCA) to generate a latent factor for ‘g’, including all of the tasks administered in the ABCD neurocognition battery.

### MRI acquisition & Image pre-processing

The ABCD MRI data were collected across 21 research sites using Siemens Prisma, GE 750 and Philips 3T scanners. Scanning protocols were harmonized across sites. Scanner ID was included in all analyses to control for any differences in image acquisition across sites and scanners. Full details of all the imaging acquisition protocols used in ABCD are outlined by Casey et al (Casey et al. 2018). Pre-processing of all MRI data for ABCD was conducted using in-house software at the Center for Multimodal Imaging and Genetics (CMIG) at University of California San Diego (UCSD) as outlined in Hagler et al (Hagler et al. 2019). Cortical surfaces were constructed from T1-weighted structural images for each subject and segmented to calculate measures of apparent cortical thickness and surface area using FreeSurfer v5.3.0(Dale et al. 1999; Fischl et al. 1999, 2004; Fischl and Dale 2000; Jovicich et al. 2006). Cortical maps were smoothed using a Gaussian kernel of 20 mm full-width half maximum (FWHM) and mapped into standardized spherical atlas space. Vertexwise data for all subjects for each morphometric measurement were concatenated into matrices in MATLAB R2017a and entered into general linear models for statistical analysis using custom written code. More details can be found in the Supplementary Materials.

## Statistical Analysis

### Behavioral analysis

All behavioral variables were standardized (z-scored) prior to analysis. Linear mixed effects models were used to residualize all behavioral variables for age, sex and a random effect of family ID prior to pairwise analysis. Partial pairwise Pearson correlations were conducted to determine associations between all of the cognitive tasks. This analysis was then repeated additionally residualizing all behavioral variables by race/ethnicity, household income and highest parental education to determine the impact of controlling for these sociodemographic factors on the pairwise associations. We additionally ran a Principal Components Analysis (PCA) across all single cognitive tasks. A scree plot of eigenvalues demonstrated that the first unrotated principal component (PC1) explained substantially more variance across the tasks than the subsequent components. The loadings of each single task with PC1 are shown in supplementary figure 1 alongside the scree plot. PC1 encompasses the shared variance across cognitive tasks, which is often described in the literature as an estimate of ‘g’: general cognitive ability. This was also included in the correlation matrices.

### Vertex-wise effect size maps

Vertex-wise data were standardized (z-scored) prior to analysis. We applied a general linear model (GLM) univariately at each vertex (n=1284) associating a given behavior from a set of covariates (age, sex, scanner ID, race/ethnicity, household income and parental education) and the vertexwise morphology data. Mass univariate standardized beta surface maps were created showing the vertex-wise associations for each analysis. Vertexwise effect size maps were calculated for all of the thirteen cognitive measures predicted by relative CSA (controlling for total CSA) and relative CTH (controlling for mean CTH) separately resulting in N=26 independent vertex-wise analyses. Models determining the association between CSA and behavior included total CSA (sum of CSA across vertices) as an additional predictor and models analyzing CTH included the mean CTH across vertices. These analyses were repeated with and without including race/ethnicity, income and parental education covariates.

### Determining the significance of the effect size maps using an omnibus test

In order to determine whether there was a significant association between vertex-wise cortical morphology (controlling for global measures) and cognitive task performance, we employed a permutation test using a univariate statistic (min-p) and three multivariate statistics (MOSTest, Stouffer and Fisher). Using the least-squares estimates *β_ν_* from each GLM at each vertex, we computed the *V x* 1 vector of Wald statistics ***z*** = (*z*,…, *z_v_*)’ across the whole cortex and accompanying p-values: *p* = 2*normcdf* (—|*z*|). Using these values, we calculated 4 different test statistics for the permutation test outlined in Table 1.

The univariate statistic, min-P was defined as the most significant (smallest) p-value across the cortex. This is a commonly used neuroimaging omnibus test, but does not take into account the distributed nature of many brain-behavior associations. The multivariate omnibus test (MOSTest)(Shadrin et al. 2020; van der Meer et al. 2020) is a more pertinent omnibus test as it determines the significance of the whole pattern of effects across the cortex taking into account the covariance structure across vertices. *χ*^2^_*MOST*_ was calculated as the estimated squared Mahalanobis norm of ***z***. The Fisher and Stouffer methods are additional alternative multivariate methods for aggregating effects across the cortex, but do not take into account the covariance across vertices. These are likely less optimal statistics, but do not rely on the regularization of a high dimensional matrix, therefore were included to show convergence of results across multiple methods.

**Table 1.**
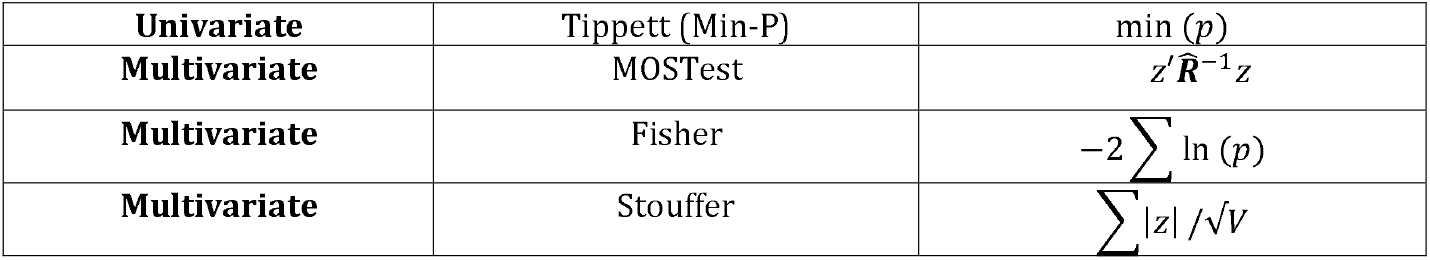
Univariate and multivariate test statistics measured.

To correctly estimate the null distribution from permutations, subject labels were shuffled according to exchangeability blocks (EBs) defined based on the family structure within ABCD (Winkler et al. 2015). This permutation procedure was used to determine the distribution of each test statistic under the global null hypothesis H_0_. We rejected H_0_ if the observed test statistic was greater than the value of the permuted test statistic at the critical threshold corresponding to an alpha level of 0.0038 (0.05 corrected for the 13 cognitive tests analyzed).

### Comparison across associations maps

To determine the shared variance between the effect size maps for the fluid (F) and crystallized (C) composite scores for each imaging phenotype, additional maps were created controlling for the other composite measure by including that measure as an additional covariate within the design matrix. This produced a vertexwise standardized beta effect size surface map for F independent of the association between cortical morphology and C (F_C_) and C independent of F (C_F_). Surface maps of the difference between the vertexwise beta coefficients as well as vertexwise Pearson correlation coefficients were calculated for the following contrasts: F – C, F – F_C_ and C – C_F_. The magnitude of effects across the cortex for each map was calculated using the following formula: 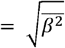. In order to determine the relative size of the effects across the difference maps compared to the individual maps we calculated a root mean square (RMS) ratio using the following formula: 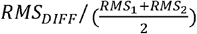, with *RMS_DIFF_* representing the RMS of the difference map and RMS1 and RMS2 as the RMS for each individual map used to produce the difference.

### Quantifying the magnitude of the association between brain structure and cognition using a mass univariate polyvertex score (PVS_U_)

PVS_U_ were calculated for the fluid and crystallized composite scores to quantify the behavioral variance explained by the vertex-wise cortical morphology. The method used here was the mass univariate PVS_U_ method outlined in detail by Zhao and colleagues(Zhao et al. 2020). In contrast to the Bayesian-PVS highlighted in their paper, the mass univariate PVS_U_ method does not take into account the covariance structure of the imaging phenotype across vertices. For vertexwise structural imaging measures the Bayesian-PVS does not increase predictive power (unlike for functional MRI), therefore, for parsimony, the mass-univariate method was used here. PVS_U_ were calculated within a 10-fold cross validation framework such that each model was trained on 90% of the sample and scores then calculated by predicting the behavior using the estimated model in the remaining 10% until a PVS_U_ was calculated for each subject. All behavioral and imaging data were pre-residualized using the covariates of no interest within each cross-validation fold prior to model estimation. PVS_U_ were calculated with and without controlling for sociodemographic variables. Global CSA and CTH measures specific to each modality were included in the pre-residualization as covariates. This allowed us to determine the unique association between relative cortical morphology and cognition and compare this to the predictive power of global measures. The association between the relative imaging phenotype and behavior across the whole sample was calculated as the squared correlation (R^2^) between the observed behavior and the predicted behavior (the PVS_U_).

In order to explore the proportion of shared variability in cognitive performance explained by brain structure and the sociodemographic variables, we generated several linear models with differing predictors. These models were generated separately for each imaging modality (when these were included) and with either the fluid or crystallized scores as the dependent variable. Each model was trained on 90% of the sample and tested in a 10% hold out set within a 10-fold cross validation framework to produce a robust, out-of-sample R^2^. The models are outlined in figure 7.

Additional models were run to estimate the out of sample variance in PC1 predicted by the relative and global imaging variables as well as the fluid and crystallized PVS_U_ in order to understand the overlap in variance across these behavioral measures. Outside of the cross-validation framework, partial correlation coefficients were estimated across the whole sample by pre-residualising both the dependent and independent variables by the covariates of no interest and/or sociodemographic variables and then correlating the residuals. All out-of-sample R^2^ estimates and partial correlation coefficients can be found in Supplementary Table 3.

## RESULTS

### Behavioral data

Figure 1A displays pairwise Pearson correlation coefficients describing the phenotypic relationship between all of the cognitive tasks measured at baseline in the ABCD study, the composite scores and the latent g-factor (PC1) controlling for age, sex and a random effect of family ID. As expected, performance across all tasks was positively correlated. Reading recognition and picture vocabulary performance were most highly correlated with each other (out of the single task measures; r=0.45). These scores were averaged to produce the crystallized composite score, therefore show a high correlation with this measure (r=0.82-0.87). The Toolbox measures used to produce the fluid composite score (flanker, dimensional card sorting, pattern processing speed, picture sequence and list working memory) were less correlated with each other (r=0.13-0.39), therefore showed slightly lower correlations with the fluid composite measure compared to the tasks contributing to the crystallized composite score (r=0.59-0.68). The total composite score (mean of fluid and crystallized) was more highly correlated with the fluid measure (r=0.90) compared to the crystallized measure (r=0.73). The matrix reasoning task showed similar correlations across the composite scores (r=0.36-0.43), which were lower than expected given that matrix reasoning is often used as a measure of fluid intelligence. As expected, the RAVLT showed the highest correlations with the verbal and memory related tasks from the NIH Toolbox. The LMT showed low correlations across all the cognitive tasks (r=0.12-0.22). PC1 derived from all single cognitive tasks showed a strong correlation with the Total Composite score from the NIH Toolbox as expected (r=0.95) and similar correlations with all of the other tasks as the Total Composite score.

**Figure 1.**
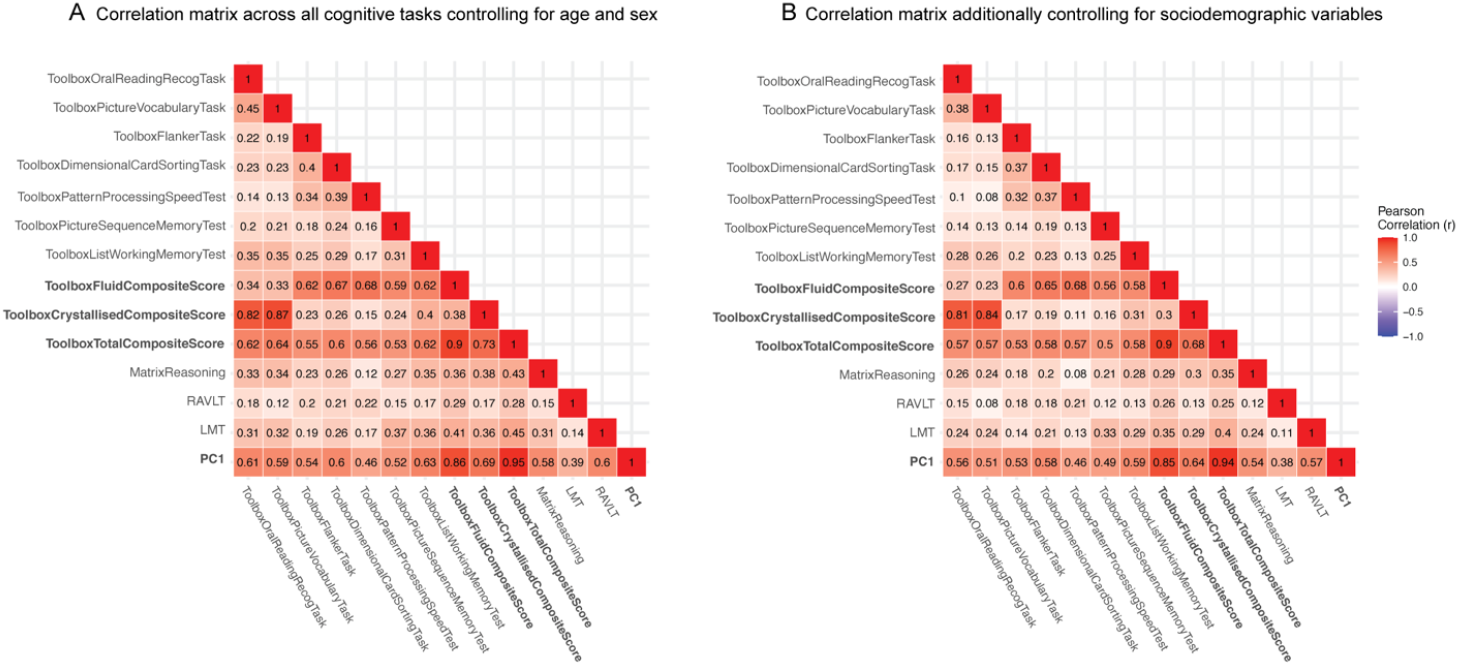
Phenotypic correlations between all of the cognitive measures at baseline in ABCD. A) Partial Pearson correlation coefficients for all cognitive tasks, composite scores and the latent factor (PC1) controlling for age, sex and a random effect of family. B) Correlations after additionally controlling for sociodemographic covariates (race/ethnicity, household income, parental education). Cognitive performance was pre-residualized for all covariates and then correlated using a Pearson correlation. Darker colors indicate a higher correlation coefficient. The composite scores and PC1 are highlighted in bold. The other cognitive measures are single tasks. Correlation coefficients among the single task measures decreased after controlling for the sociodemographic variables demonstrating that these additional measures explained a proportion of the variance in cognitive performance among these tasks.

Figure 1B shows the same correlation matrix after additionally pre-residualizing all of the cognitive measures for sociodemographic measures (race/ethnicity, household income and parental education) in order to show the partial phenotypic correlations after controlling for these confounding factors. There was a slight decrease across the correlation coefficients; however, the overall pattern of these relationships remained consistent.

### Determining the association between the regionalization of cortical morphology and cognition

We used a multivariate statistical omnibus test (MOSTest)(Shadrin et al. 2020; van der Meer et al. 2020) to measure the association between the regionalization of CSA and CTH and individual differences in cognitive performance on the fluid and crystallized composite scores from the NIH Toolbox. This multivariate approach aggregates effects across the entire cortex and therefore is better able to detect associations that are distributed compared to a standard univariate neuroimaging omnibus test that assumes effects are sparse and localized. The MOSTest also takes into account the covariance across the brain. This statistical procedure implements a permutation test with 10,000 permutations to determine statistical significance. For all associations the observed multivariate statistic fell beyond the null distribution of permuted statistics (p<0.0001; Table 2). To calculate a more precise p-value for the associations we extrapolated beyond the null distribution (see Supplementary Methods). All associations were also significant when using more widely used univariate omnibus statistics (Min-P; p<0.0005), but were smaller in magnitude than when using the MOSTest. This suggests that the distributed signal across the cortex may be important for predicting cognitive performance. The regionalization of CTH showed greater associations with cognitive performance than CSA. All of these analyses controlled for all sociodemographic factors (race/ethnicity, household income and parental education) as well as the global parameter (total CSA or mean CTH), age, sex and scanner ID. In addition, we measured all associations using two alternative multivariate statistics, Stouffer and Fisher, to determine method invariance of these brain-behavior associations. All of these methods converged (supplementary figure 2).

**Table 2.** Cortical morphology was significantly associated with both fluid and crystallized composite scores. Permuted p-values demonstrate statistical significance based on the rank of the observed statistic in the distribution of permuted statistics. For all associations both the observed univariate (min-P) and multivariate (MOST) test statistics were in the extreme of the respective null distributions, therefore we rejected the null hypothesis that the observed statistics came from the empirical null distribution. Extrapolated p-values provide an estimate of the likelihood of the observed statistics beyond the range that can be directly estimated from the permutations. The magnitude of effects was larger using the multivariate test statistic and for associations between cognitive performance and CTH (controlling for mean CTH).

|  | *Permuted P-values* |  | *Extrapolated P-values* |  |
| --- | --- | --- | --- | --- |
|  | Min-P | MOST | Min-P | MOST |
| *CSA ~ fluid score + covariates* | 5.00E-04 | 1.00E-04 | 1.29E-04 | 2.93E-10 |
| *CSA ~ crystallized score + covariates* | 1.00E-04 | 1.00E-04 | 9.34E-06 | 1.53E-07 |
| *CTH ~ fluid composite score + covariates* | 1.00E-04 | 1.00E-04 | 3.96E-05 | 5.59E-19 |
| *CTH ~ crystallized score + covariates* | 2.00E-04 | 1.00E-04 | 4.13E-08 | 8.71E-18 |

### Distinct patterns of association between the regionalization of CSA and the fluid and crystallized composite scores

After establishing significant associations between cortical morphology and cognitive performance, we aimed to measure the similarity across the estimated effect size maps for these associations. Interestingly, on visualizing the estimated effect size maps of the associations between the regionalization of CSA (controlling for total CSA) and the fluid and crystallized composite scores, we saw a unique structural pattern of association for each composite score (Fig 2A&B). Indeed, similarity maps demonstrated very little overlap between these vertexwise associations (supplementary figure 4A-C). Figure 2C shows the difference between the beta coefficients for the fluid (F) and crystallized (C) surface maps. To quantify the magnitude of these vertex-wise differences relative to the original associations, we calculated a ratio of the variance (root mean squared) of the beta coefficient differences divided by the variance (root mean squared) of the average beta coefficients across the F and C estimated effect size maps (RMS ratio for F – C = 1.21).

**Figure 2.**
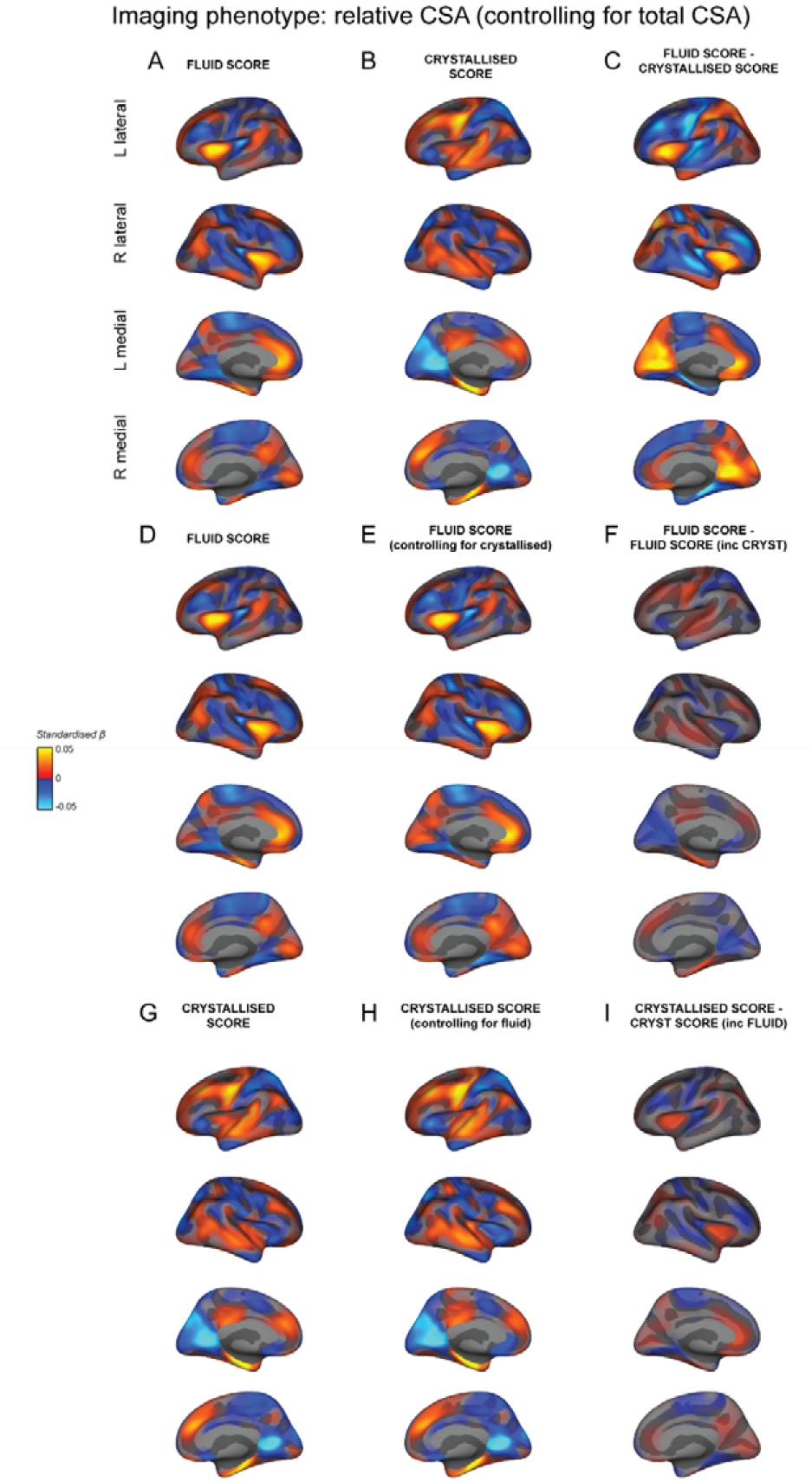
Distinct estimated effect size maps of association between the regionalization of CSA and fluid and crystallized composite scores. All maps display the vertex-wise mass univariate standardized beta coefficients unthresholded for each association. A+D) The association between relative CSA and the fluid composite score and (B+G) the crystallized composite score. C) The difference in standardized beta coefficients between A+B. E) The association between relative CSA and the fluid composite score controlling for the crystallized composite score. F) The difference in standardized beta coefficients between D+E. H) The association between relative CSA and the crystallized composite score controlling for the fluid composite score. I) The difference in standardized beta coefficients between G+H. The estimated effect size maps showing the association between the regionalization of CSA and the fluid and crystallized composite scores have distinct patterns. These behavioral measures show very little overlapping variance with the regionalization of CSA.

In order to determine the unique pattern of association between the relative configuration of CSA and the fluid and crystallized composite scores, we generated a vertex-wise association map between the regionalization of CSA and the fluid composite score controlling for the crystallized composite score (F_C_; figure 2E), and the crystallized composite score controlling for the fluid composite score (C_F_; figure 2H). The pattern of associations was almost identical to that obtained when the respective composite score was not included as a covariate in the model. Indeed, the surface map of the vertex-wise *differences* between the associations for each composite score and that composite score controlling for the other were very small in magnitude compared to the original F – C effect size maps (figure 2F,I; RMS ratio for C – C_F_ = 0.27; RMS ratio for F – F_C_ = 0.38).

Moreover, the vertex-wise correlation between the estimated beta coefficients for F vs _C_ was high (r=0.92) as was the correlation between the beta coefficients for C vs C_F_ (r=0.95). In contrast, the vertex-wise correlation between the beta coefficients for F vs C was much lower (r=0.30). This further implies that there was minimal shared variance between the associations for the fluid and crystallized scores and the regionalization of CSA. The minimal overlap in the cortical configuration associated with these measures supports that the configurations of relative CSA associated with these measures were relatively distinct.

### Distinct patterns of association between the regionalization of CTH and the fluid and crystallized composite scores

Estimated effect size maps showing the vertex-wise associations between the relative configuration of CTH and the fluid and crystallized composite scores are shown in figure 3A&B. The structural pattern of association between these composite scores and CTH were more similar than with CSA, particularly on the medial surface and similarity maps highlighting overlap in (same direction) associations showed a moderate degree of overlap (supplementary figure 4D-F). However, there were key regions with distinct differences in relative CTH associations between the fluid and crystallized composite scores. Figure 3C shows the difference between the beta coefficients for the F and C surface maps, which were relatively large compared to the original estimated effect sizes (RMS ratio for F – C = 0.92). In order to determine the unique pattern of association between the regionalization of CTH and F and C, we generated a vertex-wise map of the association between the regionalization of CTH and F_C_ (figure 3E) and CTH and C_F_ (figure 3H). As with CSA, the pattern of associations was almost identical to when the respective intelligence measure was not included in the model. Indeed, the estimated effect size map of the vertex-wise differences between the associations for each composite score and each composite score controlling for the other were very small in magnitude (figure 3F,I; RMS ratio for C – C_F_ = 0.31; RMS ratio for F – F_C_ = 0.30). The vertexwise correlations between the map of associations for each composite score and that composite score controlling for the other were high (F vs F_C_: r=0.94; C vs C_F_: r=0.95); whereas the vertex-wise correlation between the maps of association for the fluid and crystallized composite scores was much lower (F vs C: r=0.40). The minimal overlap in the cortical configurations associated with these measures supports that the configurations of CTH associated with these measures were relatively distinct.

**Figure 3.**
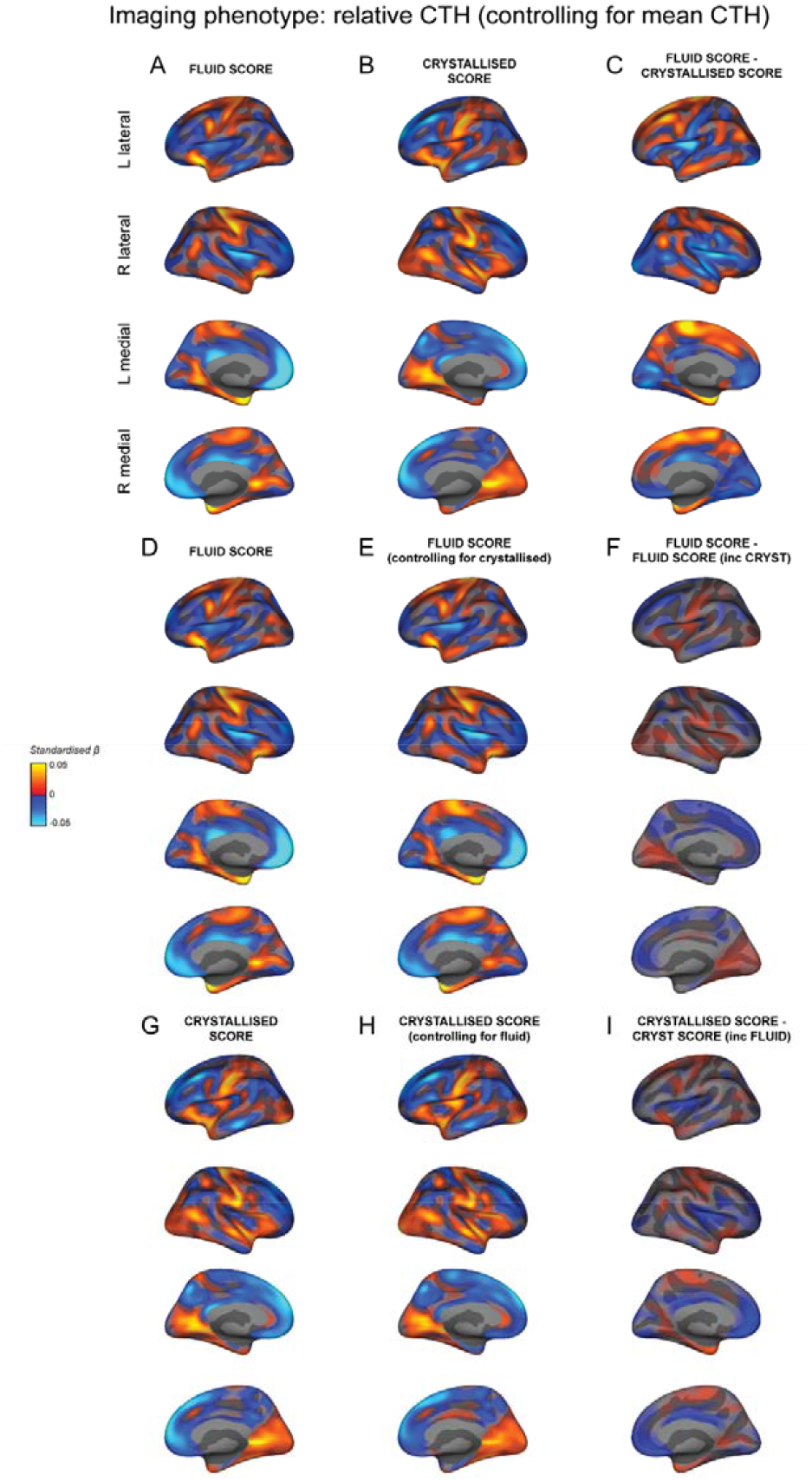
Distinct estimated effect size maps of association between the regionalization of CTH and fluid and crystallized composite scores. All maps display the vertexwise mass univariate standardized beta coefficients unthresholded for each association. A+D) The association between relative CTH and the fluid composite score and (B+G) the crystallized composite score. C) The difference in standardized beta coefficients between A+B. E) The association between relative CTH and the fluid composite score controlling for the crystallized composite score. F) The difference in standardized beta coefficients between D+E. H) The association between relative CTH and the crystallized composite score controlling for fluid composite score. I) The difference in standardized beta coefficients between G+H. The estimated effect size maps showing the association between the regionalization of CTH and the fluid and crystallized composite scores have distinct patterns. These behavioral measures show veiy little overlapping variance with the regionalization of CTH.

### Distinct patterns of association for the regionalization of CTH and CSA across cognitive tasks

Visualizing the effect size maps between each cortical measure and each cognitive task revealed relatively distinct patterns of association across the cognitive tasks (figure 4). This was further supported by the low vertex-wise correlations across these associations (figure 5). The estimated beta coefficients for the regionalization of CTH and cognition were more correlated across tasks than for the regionalization of CSA and cognition. Effect size maps were more similar for tasks using similar underlying cognitive processes and the composite measures appeared to reflect mixtures of the patterns of association for each of the tasks that were averaged to produce the composite scores. Estimated effect size maps for each cortical measure and each cognitive task and composite score without controlling for the sociodemographic variables of race/ethnicity, household income and parental education were more highly correlated (figure 5C&D). This shows that variance in the sociodemographic factors is shared with the cognitive and structural measures. The association maps for PC1 and the Total Composite Score from the NIH Toolbox were remarkably similar further validating the Total Composite Score as a measure of general cognitive ability.

**Figure 4.**
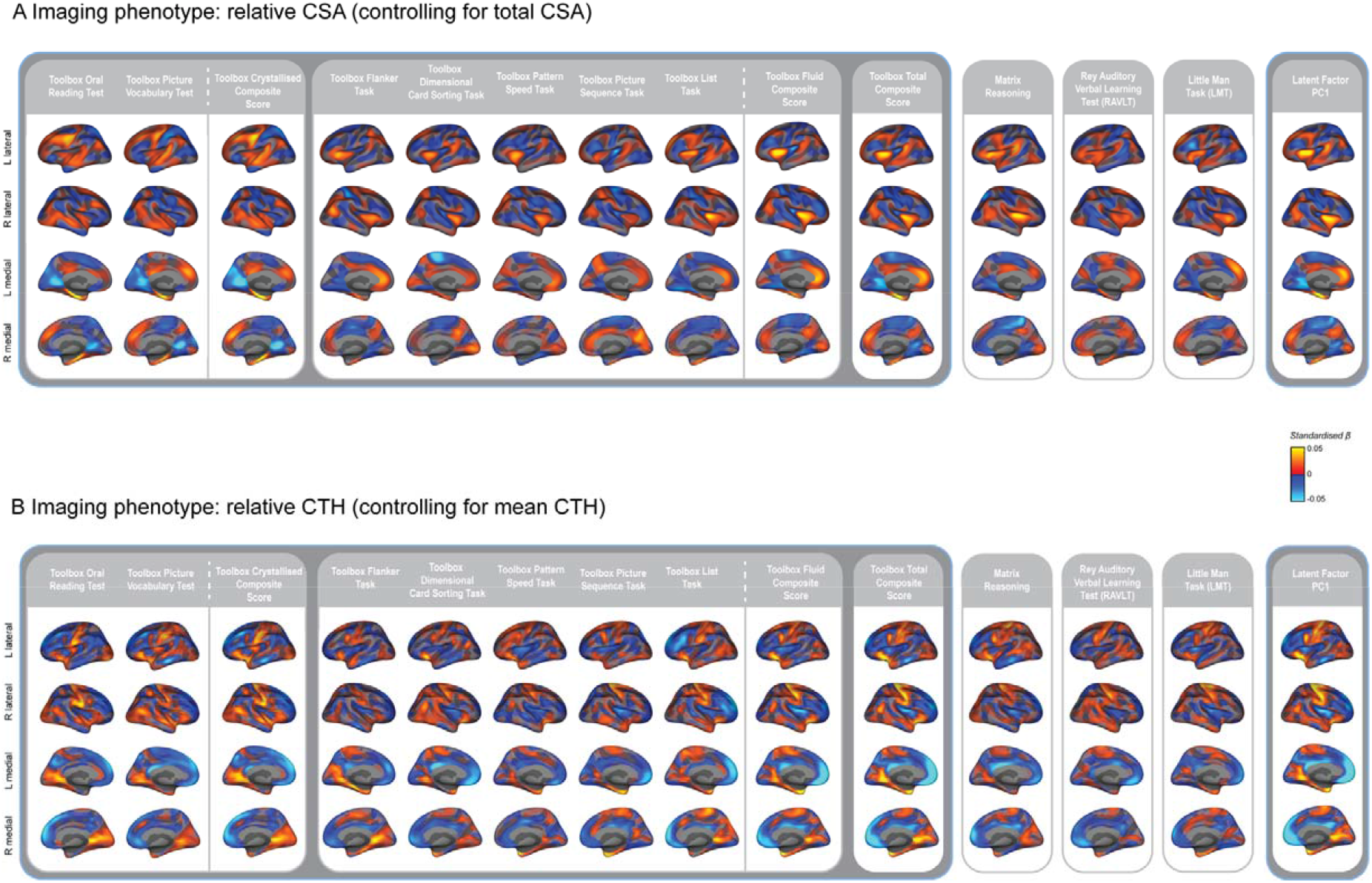
Estimated effect size maps showing the mass univariate standardized beta coefficients for the association between each cognitive task and (A) the regionalization of CSA and (B) the regionalization of CTH. Performance on all of the cognitive tasks and the composite scores showed significant associations with the regionalization of cortical morphology across the whole cortical surface according to the MOSTest as determined in Palmer et al (2019).

**Figure 5.**
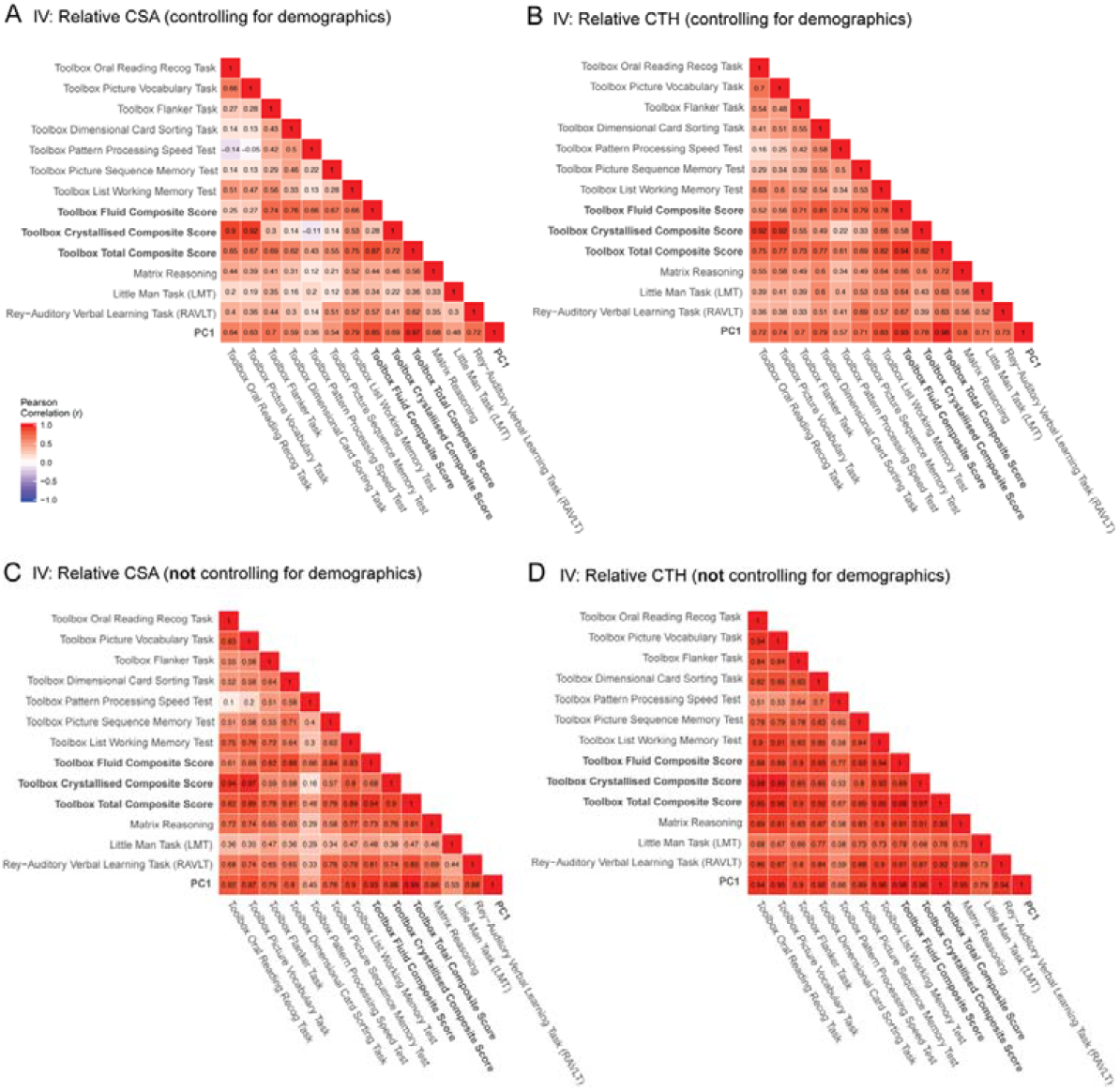
Similarity of regionalization association patterns across cognitive tasks is modulated by demographic factors. Pairwise vertex-wise correlations between the estimated beta coefficients across all cognitive tasks, the composite scores and **PC1** predicted by A) the regionalization of CSA and B) the regionalization of CTH controlling for the sociodemographic variables of race/ethnicity, household income and parental education. C&D) Pairwise vertex-wise correlations for all of the cognitive tasks and C) the regionalization of CSA and D) the regionalization of CTH <u>not</u> controlling for the specified sociodemographic variables (but controlling for age, sex and scanner). The correlation amongst associations was larger when these demographic variables were not included in the GLMs used to produce the estimated effect size maps.

### Association maps when not controlling for sociodemographic variables

We computed the estimated effect size maps for the F and C composites scores associated with the regionalization of CSA (figure 6A&B) and CTH (figure 6D&E) without controlling for the sociodemographic variables (but controlling for age, sex and scanner). For both morphology measures, the vertex-wise correlation between the F and C maps increased (CSA: r=0.69; CTH: r=0.89). However, the F-C difference maps were remarkably similar to when the sociodemographic variables were controlled for as shown by a vertex-wise correlation between the F-C difference maps with and without controlling for sociodemographic factors (CSA: r=0.96; CTH: r=0.94). This strongly suggests that these confounding variables in aggregate index common shared variance between cortical morphology and cognition across domains that is not specific to a particular task performance. Estimated effect size maps for the association between the regionalization of CSA and CTH and cognitive performance across tasks without controlling for sociodemographic factors can be found in supplementary figure 5.

**Figure 6.**
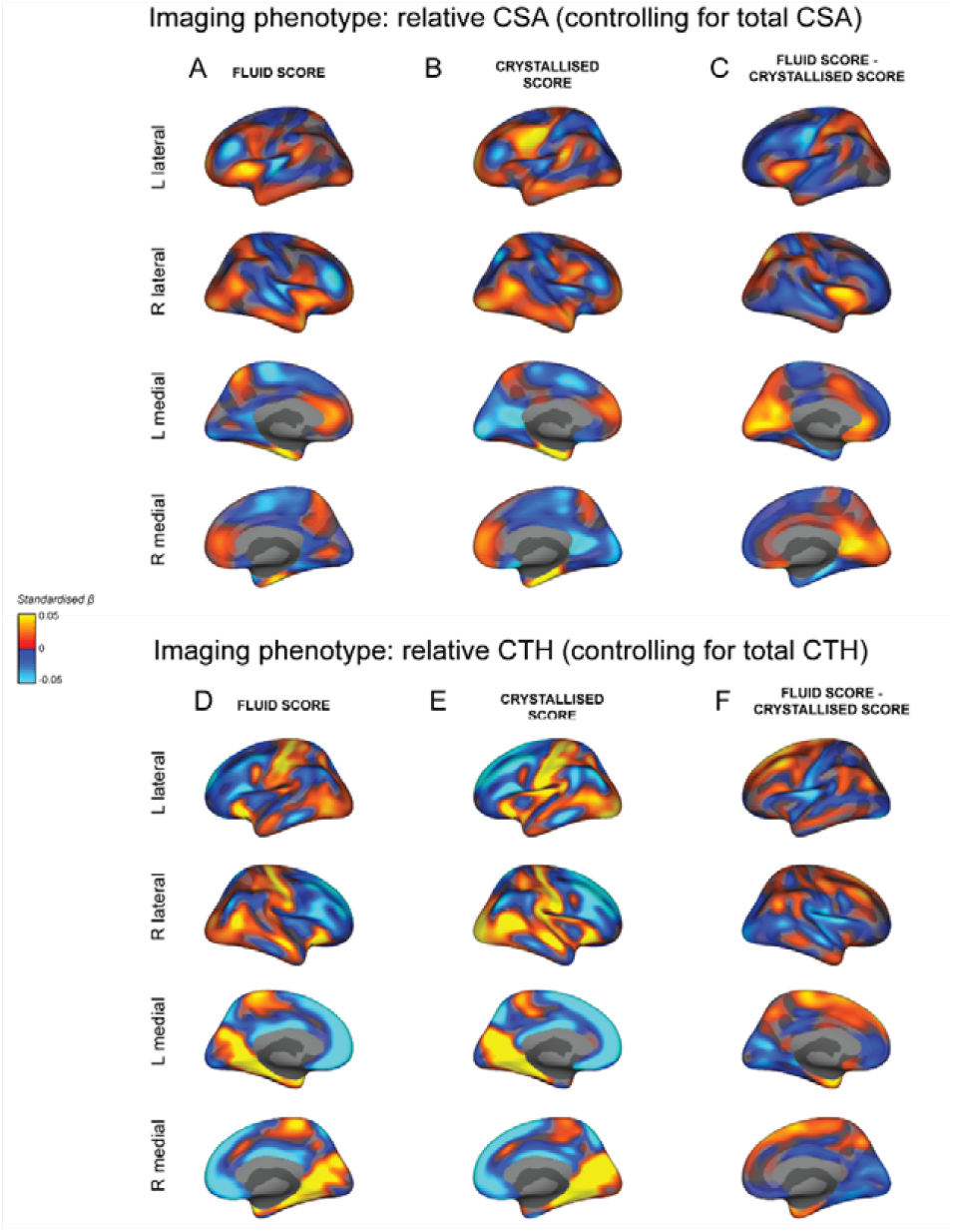
Similar estimated effect size maps for the association between the regionalization of cortical morphology and the fluid and crystallized composite scores when <u>not</u> controlling for sociodemographic factors. All maps display the vertex-wise mass univariate standardized beta coefficients unthresholded for each association. The top part of the figure shows the association between the regionalization of CSA and the fluid composite score (A) and the crystallized composite score (B). C) The difference in standardized beta coefficients between A+B. The bottom part of the figure shows the association between the regionalization of CTH and the fluid composite score (D) and the crystallized composite score (E). F) The difference in standardized beta coefficients between D+E. The demographic variables of race/ethnicity, household income and parental education were not included in the mass univariate GLMs to produce these estimated effect size maps. The exclusion of these confounding factors increased the magnitude of the estimated beta coefficients and the similarity between the maps for the fluid and crystallized composite scores. However, the difference maps (C+F) closely reflect the pattern of differences seen in figure 2C and figure 3C respectively.

### Partitioning the variance between brain structure and cognition

In order to quantify the variance in each composite score predicted by the vertex-wise imaging data for the regionalization of CSA and CTH, we calculated a mass univariate polyvertex score (PVS_U_). Much like a polygenic risk score, the PVS_U_ represents a linear weighted sum of the vertex-wise associations, which are projected from a training set to an independent hold-out set within a cross validation framework. A comparison of the subject-wise PVS_U_ with cognitive performance therefore provides a conservative, out-of-sample, lower-bound estimate of the unique variance in behavior predicted by the regionalization of cortical morphology independent of the global brain measures. When controlling for age, sex and scanner, the regionalization of CSA explained a similar proportion of the total variability in cognition compared to the regionalization of CTH (for fluid scores: total CSA = ~2.59%R^2^; CSA PVS_U_ = ~1.27%R^2^; CTH PVS_U_ =~1.88%R^2^; for crystallized scores: total CSA predicted ~7.15%R^2^; CSA PVS_U_ predicted ~2.39%R^2^; CTH PVS_U_ = ~3.85%R^2^; figure 7). Mean CTH was not predictive of cognitive performance. Total CSA predicted more variance in crystallized scores than fluid scores.

**Figure 7.**
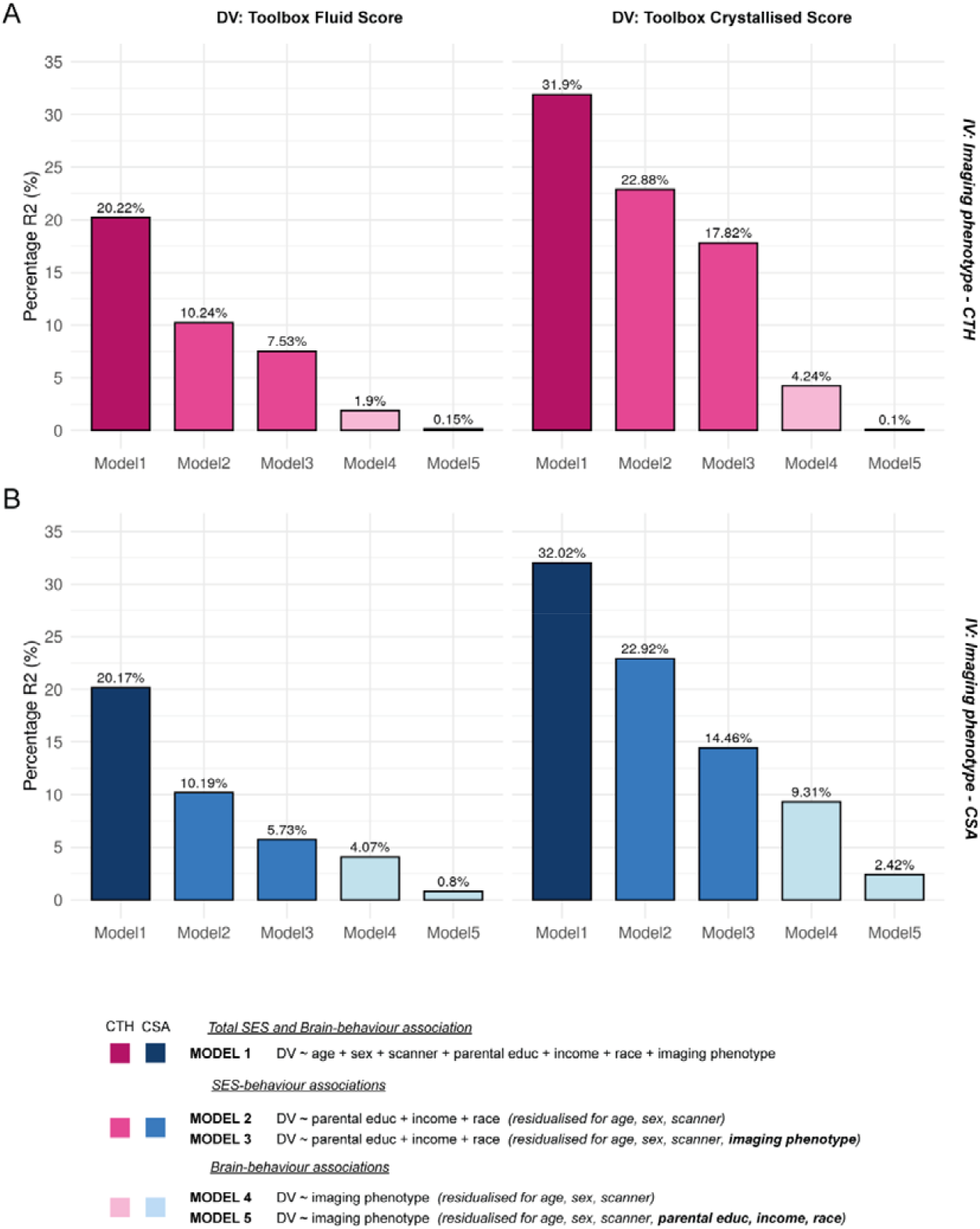
Controlling for sociodemographic factors reduces the variance in cognitive performance predicted by cortical morphology. These bar charts show the percentage R^2^ in the fluid (left column) and crystallized (right column) composite scores predicted by several models including sociodemographic and brain measures. Model 1 shows the total variance in behavior explained by the full model including the covariates of no interest (age, sex and scanner), the sociodemographic variables (parental education, household income and race/ethnicity) and either the regionalization of CTH and mean CTH (A; pink) or the regionalization of CSA and total CSA (B; blue). Model 2 shows the unique variance in behavior explained by the sociodemographic factors after pre-residualizing both the dependent (DV) and independent (IV) variables for the covariates of no interest. Additionally, pre-residualizing for the imaging phenotypes (model 3) lead to a reduction in this R^2^. Conversely, model 4 shows the *unique* variance in behavior explained by the structural measures (global and regionalization) after pre-residualizing for the covariates of no interest only. Additionally, pre-residualizing for the sociodemographic factors (model 5) lead to a large decrease in the variance explained by these structural measures, again showing the shared variance between brain structure, cognition and sociodemographic factors. Controlling for the sociodemographic factors reduced the variance explained in the composite scores by ~4-5 fold for CSA and ~4-8 fold for CTH.

Without controlling for brain structure, but whilst controlling for age, sex and scanner, the sociodemographic factors collectively predicted more variance in crystallized (~23%R^2^) compared to fluid scores (~10%R^2^). Across imaging measures, the sociodemographic variables accounted for ~70-90% of the variation in fluid and crystallized scores that was predicted by brain structure. CTH measures (mean CTH and CTH PVS_U_) explained ~22% of the association between the sociodemographic variables and the crystallized score and ~27% for the fluid score, whilst CSA (total CSA and CSA PVS_U_) explained ~37% of the association between the sociodemographic variables and the crystallized score and ~44% for the fluid score. This shows that the relationship between brain structure and cognition was strongly related to the sociodemographic factors of race/ethnicity, parental education and household income.

We additionally estimated the variance in PC1 predicted by regional CSA and CTH measures using the PC1 PVS_U_ when controlling and not controlling for sociodemographic variables. Cross-validated R^2^ estimates were greatly reduced when controlling for sociodemographic variables (Supplementary Table 3). When controlling for sociodemographic variables in particular, the fluid and crystallized PVS_U_ predicted less variance in the behavior they were not trained on highlighting key differences in the associated regionalization patterns. Moreover, a model predicting PC1 from both the fluid and crystallized PVS_U_ (for CSA and CTH independently) explained a similar proportion of the variance to the sum of R^2^ across separate models with the fluid and crystallized PVS_U_ predicting PC1. This was less clear for CTH due to the greater similarity in relative association patterns. When not controlling for sociodemographic variables, the overlap in the variance in PC1 explained by the fluid and crystallized PVS_U_ was much greater. This emphasizes that much of the shared variance between cognitive tasks and associated regionalization patterns is associated with sociodemographic factors.

## DISCUSSION

In this study, we have shown that the regionalization of cortical morphology, independent of global brain measures, was significantly associated with individual differences in cognitive performance in a large sample of 9-11-year-old children (N=10,145). Moreover, we showed that individual differences in composite scores of fluid and crystallized performance were associated with distinct maps of relative cortical areal expansion and apparent thickness. This suggests that regional cortical architecture relates differently to these two types of cognitive performance variability. The relatively distinct association maps for the single tasks used to generate these composite measures provide evidence that patterns of cortical morphology differ among individuals and can explain individual variability across different cognitive processes. These patterns do not appear to represent a single, neural construct that could explain the positive manifold. Furthermore, we have shown that sociodemographic diversity impacts the association between cortical morphology and cognition similarly across cognitive domains, which suggests these factors are likely explaining confounding variance to the relationship between brain structure and cognition. Since the treatment of sociodemographic variables in these models led to substantial differences in the magnitude of effects of interest, users of ABCD data should select covariates carefully and discuss any sociodemographic factors potentially related to both explanatory variables and outcomes.

### Relative cortical configuration is relevant for understanding individual variability in cognition

There is a high degree of individual variability in the regionalization of the cortex, however the extent to which that can explain individual variability in behavior is not well understood. A number of studies have highlighted associations between regional differences in cortical morphology, controlling for global imaging measures, and cognition; however, they have lacked the power to measure associations based on the entire configuration of cortical morphology(Vuoksimaa et al. 2016). Individual differences in the regionalization of CSA and CTH have been associated with distinct, continuous gradients of genetic influences across the cortex (Chen et al. 2011; Chen, Fiecas, et al. 2013; Chen, Gutierrez, et al. 2013) that coincide with the gene expression patterns that dictate specialization of the neocortex during embryonic development (O’Leary et al. 2007; Rakic et al. 2009). Individual differences in this molecular signaling could therefore lead to subtle alterations in this cortical configuration, which could lead to variability in behavior. Here we have used a multivariate statistical approach to measure the significance of distributed effects, which reflect the graded and distributed nature of the biology of regionalization.

We have shown that individual variability in cortical regionalization is robustly associated with individual variability in performance across multiple cognitive tasks. For both morphology measures, there was a clear pattern of positive and negative associations with cognitive performance, which suggests that, at this developmental stage, *relative* differences in the size and thickness of cortical regions mirror individual difference variability in cognition. Although we have corrected for age in all of our models, we are only measuring associations at a single time-point, therefore phase differences in cortical maturation may be contributing to variability in regionalization. Nevertheless, the regionalization of CSA predicted additional unique variance in cognition not explained by total CSA. By partitioning the variance in this way, we can determine the relative importance of these components of CSA for cognition. This appeared to differ depending on the cognitive task, with more language related measures having a greater association with total CSA compared to the regionalization of CSA. Comparatively the proportion of variance in behavior explained by regional CTH was relatively similar across the different measures. Consistent with previous studies (Vuoksimaa et al. 2015, 2016; Schmitt et al. 2019; Grasby et al. 2020), mean CTH did not predict cognitive performance across any measures.

### Distinct associations between cortical morphology and different cognitive domains

The estimated effect size maps between the regionalization of CSA and the crystallized composite score (comprised of reading and vocabulary measures) showed that greater areal expansion in regions previously associated with language functions, such as the left temporal and middle frontal regions(Brown et al. 2001; Hickok and Poeppel 2007; Martin et al. 2015), was associated with higher crystallized scores; whereas, greater areal expansion in a different set of regions, previously implicated in cognitive control mechanisms required for the fluid tasks, such as the anterior cingulate and insula cortices(Botvinick et al. 2004; Taylor et al. 2009; Menon and Uddin 2010; Fjell et al. 2012; Curley et al. 2018), was associated with higher fluid scores. The patterns of apparent CTH associated with these measures were more similar, however there were key differences in lateral frontal, insula, superior medial and entorhinal cortices.

Interestingly, the areas most strongly associated with the total composite score were a combination of those most strongly associated with either the fluid or crystallized composite scores and were very similar to the association maps for PC1. Consistent with previous literature, the CSA and CTH maps for our measures of general cognitive ability were very similar to those observed by Vuoksimaa et al (2016) despite the difference in age between the two samples. In particular, negative associations with relative CTH and general cognitive ability were found in medial and lateral frontal cortex and positive associations around the central gyrus; and for relative CSA we observed similar positive associations across the temporal lobe, medial frontal and anterior cingulate cortices. Our results also mirror some associations in highly evolutionary and developmentally expanded regions, namely the frontal cortex, which have been shown previously to associate with general cognitive ability (Fjell et al. 2015; Reardon et al. 2018). The maps of association between both the regionalization of CSA and CTH and the total composite score (and PC1) represent a mixture of the fluid and crystallized associations. Indeed, the same can be seen for the fluid and crystallized composite score maps, which appear to represent a mixture of the association maps for each of the individual tasks that contributed to those scores. This is reflected within the correlation matrix of the vertex-wise associations (figure 5A&B). This observation has implications for theories of intellectual development.

### Implications for theories of intellectual development

Factor analyses of cognitive measures frequently reveal a higher order general latent factor ‘g’ that can explain (statistically) individual differences in cognitive performance. However, this observation alone does not necessarily reveal anything about neural underpinnings of ‘g’. There is a pervasive view within society that ‘g’ represents a single causal underlying trait that influences performance across multiple domains. GWAS have begun to reveal some genetic loci associated with general cognitive ability that exhibit pleiotropy with structural brain phenotypes, which point to biological pathways from our genes to ‘g’ via brain phenotypes (although the direction of causality is not clear)(Davies et al. 2018; Savage et al. 2018; Grasby et al. 2020). However, this does not necessarily suggest that there is a unitary pathway underlying the development of cognitive function across individuals. Indeed, polygenic scores for educational attainment and general cognitive ability differentially predict cognitive performance in different domains (Elliott et al. 2019; Loughnan et al. 2019; Mitchell et al. 2020). Here we do not find that there is a single pattern of regionalization that could represent a causal neuroanatomical substrate underlying ‘g’. Indeed, we find that, on average, individuals with one regionalization pattern perform better on more language related tasks while individuals with a different regionalization pattern perform better on more executive function related tasks. Therefore, this reveals important heterogeneity amongst children with comparable levels of performance on measures of general cognitive ability. Our results do not test any causality between brain structure and behavior and do not rule out that a single or global neural phenotype underlying (or contributing to) ‘g’ could be represented within a different neural modality, such as functional connectivity or latency of neuronal signaling; however, there is a lack of supporting evidence for a dominant neural ‘g’ phenotype when measuring cortical morphology in the current study.

Total CSA was modestly associated with cognitive performance in the current study; however, before controlling for sociodemographic factors, total CSA only predicted ~7% of the variability in crystallized scores and ~2% in fluid scores. Due to the unprecedented statistical power in this study, we can infer that total CSA is unlikely to account for substantially more individual variability in cognitive scores in children of this age. Indeed, this effect is much smaller than the shared variance across cognitive measures that has been attributed to ‘g’ (~37% in the current sample) and is of a similar magnitude to that observed by Reardon et al (2018; 1%). Moreover, the relative importance of this measure for predicting individual variability in cognition differed as a function of the type of intelligence measured, which is inconsistent with this being a global mediator of ‘g’.

Our results may be more compatible with alternative models of intellectual development in which a single structural phenotype is not necessarily required to produce the latent g-factor. For example, according to the mutualism model (Van Der Maas et al. 2006, 2017), at the beginning of development, individuals start with initially uncorrelated, differentially weighted resources that underlie different cognitive processes. Over time, these resources interact in a mutually beneficial way such that high performance of one system aids the development of another system leading to the positive manifold in performance. This is an important distinction from the more conventional ‘g’ model as here individual differences in mature cognitive performance emerge from different combinations of these resources in early development, such that, for example, the same general cognitive performance score later in development could arise from high initial memory resources in one individual, but high initial processing speed in another individual. Given the development of the positive manifold throughout childhood, we would not necessarily expect to see a large influence of these initial biases on individual variability in cognitive performance at 9-10 years old. Indeed, we can see that multiple environmental and experiential factors contribute to differences in cognitive performance across individuals. It is possible that the small, dissociable variance across cognitive domains associated with cortical regionalization, after controlling for sociodemographic factors, reflects these initial biases in resource allocation. Only in tasks that disproportionately recruit specific cognitive processes can we really detect the small differences in performance across individuals related to cortical regionalization. This may explain why we see such a small association between cortical regionalization and individual variability in cognitive performance.

Given this hypothesis, it is possible that this variability in regionalization is mediated by genetic variability in cortical arealization. Indeed, the regionalization of the cortex is moderately heritable(Winkler et al. 2010; Chen et al. 2011; Eyler et al. 2011, 2012; Chen, Gutierrez, et al. 2013) and during early embryonic development, gradients of morphogens dictate the expression of transcription factors across the dorsal proliferative zone, which determines the eventual spatial location and functional specialization of cortical projection neurons(O’Leary et al. 2007; Rakic et al. 2009). Small biases in the arealization process, reflected in variability in cortical regionalization, may influence developing cognitive functions in individuals in ways that advantage some functional domains relative to others, which creates important diversity within our population. Although the current study does not measure genetic variation directly, this hypothesis supports the magnitude of the observations here. Exposure to different environments, such as access to good education, can then disproportionately enhance specific genetic effects as has previously been shown (Dickens and Flynn 2001; Harden et al. 2007; Kan et al. 2013; Loughnan et al. 2019) and contribute to individual differences in cognitive function. We have not directly tested this theory in the current paper, however our observations are in line with what we would expect based on these models. Alternative models such as sampling models and hierarchical factor models could also explain the observations here. Longitudinal, developmental studies are required to tease apart the predictions of these different models at different developmental stages.

It is unclear how heterogenous the regionalization of cortical morphology is; however, if there is a mixture distribution, as we expect from the given observations and the mutualism model, then this would also reduce the estimated effect size from the mean association maps across the full sample. Future analyses should use data driven approaches to classify individuals based on similarity in cortical architecture and determine subsequent associations with behavior. This would also elucidate the extent to which these small associations are driven by subgroups of individuals who may have occult or prodromal pathology.

### Understanding individual variability in cognition related to both brain structure and sociodemographic factors

After controlling for age, sex and scanner, but before controlling for sociodemographic variables, the proportion of variance in the composite measures of cognitive performance shared with the brain phenotypes in aggregate was ~9% for the crystallized scores and ~4% for the fluid scores. Overall, the full model including all the phenotypes together accounted statistically for ~32% of the variance in crystallized scores and ~20% of the variance in fluid scores. These results suggest that many factors relating to individual differences in cognitive test scores remain unaccounted for in these statistical models. However, the results are sufficiently powerful to constrain future hypotheses by restricting the range of plausible causal effects of well-measured variables, such as total CSA.

The regionalization of CSA and CTH both accounted for additional unique variance in cognition independent of global measures. Even after pre-residualizing for sociodemographic factors that could index cultural, environmental and other experiential effects on cognitive test scores, as well as global brain measures, the associations between the cortical regionalization phenotypes and cognitive test scores were statistically robust; but only explained ~0.1% of the residual variation. Here we have used a very conservative, out-of-sample effect size; we are therefore providing the lower bound for the estimated effect size between regionalization and cognition, which is not influenced by any confounding measures. Given the above hypothesis regarding what would be predicted from the mutualism model, it is perhaps unsurprising that we see such a small brain-behavior association, particularly as we’re measuring individual variability in typically developing children and not pathology. Here much of the individual variability in regionalization is attributable to environmental factors, such as sociodemographic variables, although gene-environment correlations may also be contributing to this shared relationship. Moreover, small effects are common-place among samples of this size where effect sizes are not inflated by sampling variability or publication bias (Dick et al. 2020). Large brain-behavior associations in smaller studies with less precision around effect size estimates should be interpreted very cautiously.

Our results show that the sociodemographic variables accounted for a substantial proportion of the shared variance between the brain phenotypes and composite measures of cognitive function. This may result from many small effects of factors indexed by these variables, such as nutrition, limited access to healthcare and education, untreated prenatal complications, systemic racism, high levels of stress or exposure to environmental toxins. These factors may affect both brain and cognitive development, however it is not clear to what extent these follow overlapping or independent pathways, and therefore, to what extent these sociodemographic factors are mediators vs confounders in the relationship between brain structure and cognition. However, the finding that these sociodemographic factors predicted similar variance across cognitive tasks, and the association maps were more similar across tasks when *not* controlling for these factors, suggests that these variables are more likely to be confounding. Only by controlling for these measures do we remove that confounding variance and reveal specific associations, which may more closely represent causal relationships between brain structure and cognition, however, importantly, we cannot infer any causality from the current analyses.

### Conclusions

The graded and distributed nature of the neurobiology underlying the regionalization of the cortex supports the importance of studying the whole configuration of cortical morphology associated with behavior using multivariate statistics and continuous pattern comparison methods. Here we have shown a robust association between cortical regionalization and cognitive performance and have highlighted key differences in the variability of cortical regionalization that can explain individual variability across different components of cognition. Using the unprecedented ABCD dataset, we now have the power to discover novel brain-behavior associations and understand individual variability in behavior among a diverse sample of participants. With future releases of the ABCD data we will aim to better understand the heterogeneity of cortical regionalisation across the sample and track how the relationship between regionalization and cognitive performance may change over time within an individual.

## Acknowledgements

The authors wish to thank the youth and families participating in the Adolescent Brain Cognitive Development (ABCD) Study and all ABCD staff involved in data collection and curation.

## REFERENCES

Akshoomoff N, Beaumont JL, Bauer PJ, Dikmen SS, Gershon RC, Mungas D, Slotkin J, Tulsky D, Weintraub S, Zelazo PD, et al. 2013. NIH toolbox cognition battery (CB): Composite scores of crystallized, fluid, and overall cognition. Monogr Soc Res Child Dev. 78:119–132.

Arendt, Nielsen J. 2005. Does education cause better health? A panel data analysis using school reforms for identification. Econ Educ Rev. 24:149–160.

Basten U, Hilger K, Fiebach CJ. 2015. Where smart brains are different: A quantitative meta-analysis of functional and structural brain imaging studies on intelligence. Intelligence. 51:10–27.

Botvinick MM, Cohen JD, Carter CS. 2004. Conflict monitoring and anterior cingulate cortex: an update. Trends Cogn Sci. 8:539–546.

Bouchard T, McGue M. 1981. Familial studies of intelligence: a review. Science (80-). 212:1055–1059.

Brouwer RM, van Soelen ILC, Swagerman SC, Schnack HG, Ehli EA, Kahn RS, Hulshoff Pol HE, Boomsma DI. 2014. Genetic associations between intelligence and cortical thickness emerge at the start of puberty. Hum Brain Mapp. 35:3760–3773.

Brown WE, Eliez S, Menon V, Rumsey JM, White CD, Reiss AL. 2001. Preliminary evidence of widespread morphological variations of the brain in dyslexia. Neurology. 56:781–783.

Burgaleta M, Johnson W, Waber DP, Colom R, Karama S. 2014. Cognitive ability changes and dynamics of cortical thickness development in healthy children and adolescents. Neuroimage. 84:810–819.

Casey BJ, Cannonier T, Conley MI, Cohen AO, Barch DM, Heitzeg MM, Soules ME, Teslovich T, Dellarco D V, Garavan H, et al. 2018. The Adolescent Brain Cognitive Development (ABCD) study: Imaging acquisition across 21 sites. Dev Cogn Neurosci. 32:43–54.

Chen C-H, Fiecas M, Gutierrez ED, Panizzon MS, Eyler LT, Vuoksimaa E, Thompson WK, Fennema-Notestine C, Hagler DJ, Jernigan TL, et al. 2013. Genetic topography of brain morphology. Proc Natl Acad Sci. 110:17089–17094.

Chen C-H, Panizzon MS, Eyler LT, Jernigan TL, Thompson W, Fennema-Notestine C, Jak AJ, Neale MC, Franz CE, Hamza S, et al. 2011. Genetic Influences on Cortical Regionalization in the Human Brain. Neuron. 72:537–544.

Chen C, Gutierrez ED, Thompson W, Panizzon MS, Terry L, Eyler LT, Fennema-notestine C, Jak AJ, Michael C, Franz CE, et al. 2013. Hierarchical Genetic Organisation of Human Cortical Surface Area. 335:1634–1636.

Colom R, Burgaleta M, Román FJ, Karama S, Álvarez-Linera J, Abad FJ, Martínez K, Quiroga MÁ, Haier RJ. 2013. Neuroanatomic overlap between intelligence and cognitive factors: Morphometry methods provide support for the key role of the frontal lobes. Neuroimage. 72:143–152.

Colom R, Haier RJ, Head K, Álvarez-Linera J, Quiroga MÁ, Shih PC, Jung RE. 2009. Gray matter correlates of fluid, crystallized, and spatial intelligence: Testing the P-FIT model. Intelligence. 37:124–135.

Compton WM, Dowling GJ, Garavan H. 2019. Ensuring the Best Use of Data. JAMA Pediatr.

Curley LB, Newman E, Thompson WK, Brown TT, Hagler DJ, Akshoomoff N, Reuter C, Dale AM, Jernigan TL, Jernigan TL. 2018. Cortical morphology of the pars opercularis and its relationship to motor-inhibitory performance in a longitudinal, developing cohort. Brain Struct Funct. 223:211–220.

Cutler DM, Lleras-Muney A, Cutler D, Lleras-Muney A. 2012. Education and Health: Insights from International Comparisons.

Dale AM, Fischl B, Sereno MI 1999. Cortical Surface-Based Analysis. Neuroimage. 9:179–194.

Davies G, Lam M, Harris SE, Trampush JW, Luciano M, Hill WD, Hagenaars SP, Ritchie SJ, Marioni RE, Fawns-Ritchie C, et al. 2018. Study of 300,486 individuals identifies 148 independent genetic loci influencing general cognitive function. Nat Commun. 9.

Deary IJ. 2012. Intelligence. Annu Rev Psychol. 63:453–482.

Dick AS, Watts AL, Heeringa S, Lopez DA, Bartsch H, Chieh Fan C, Palmer C, Reuter C, Marshall A, Haist F, et al. 2020. Meaningful Effects in the Adolescent Brain Cognitive Development Study. bioRxiv. 2020.09.01.276451.

Dickens WT, Flynn JR. 2001. Heritability estimates versus large environmental effects: the IQ paradox resolved. Psychol Rev. 108:346–369.

Duncan J, Seitz RJ, Kolodny J, Bor D, Herzog H, Ahmed A, Newell FN, Emslie H. 2000. A neural basis for general intelligence. Science (80-). 289:457–460.

Elliott ML, Belsky DW, Anderson K, Corcoran DL, Ge T, Knodt A, Prinz JA, Sugden K, Williams B, Ireland D, et al. 2019. A Polygenic Score for Higher Educational Attainment is Associated with Larger Brains. Cereb Cortex. 29:3496–3504.

Eyler LT, Chen CH, Panizzon MS, Fennema-Notestine C, Neale MC, Jak A, Jernigan TL, Fischl B, Franz CE, Lyons MJ, et al. 2012. A comparison of heritability maps of cortical surface area and thickness and the influence of adjustment for whole brain measures: A magnetic resonance imaging twin study. Twin Res Hum Genet. 15:304–314.

Eyler LT, Prom-Wormley E, Panizzon MS, Kaup AR, Fennema-Notestine C, Neale MC, Jernigan TL, Fischl B, Franz CE, Lyons MJ, et al. 2011. Genetic and environmental contributions to regional cortical surface area in humans: A magnetic resonance imaging twin study. Cereb Cortex. 21:2313–2321.

Fischl B, Dale AM. 2000. Measuring the thickness of the human cerebral cortex from magnetic resonance images. Proc Natl Acad Sci U S A. 97:11050–11055.

Fischl B, Salat DH, van der Kouwe AJW, Makris N, Ségonne F, Quinn BT, Dale AM. 2004. Sequence-independent segmentation of magnetic resonance images. Neuroimage. 23:S69–S84.

Fischl B, Sereno MI, Dale AM. 1999. Cortical Surface-Based Analysis II: Inflation, Flattening, and a Surface-Based Coordinate System.

Fjell AM, Walhovd KB, Brown TT, Kuperman JM, Chung Y, Hagler DJ, Venkatraman V, Roddey JC, Erhart M, McCabe C, et al. 2012. Multimodal imaging of the self-regulating developing brain. Proc Natl Acad Sci U S A. 109:19620–19625.

Fjell AM, Westlye LT, Amlien I, Tamnes CK, Grydeland H, Engvig A, Espeseth T, Reinvang I, Lundervold AJ, Lundervold A, et al. 2015. High-Expanding Cortical Regions in Human Development and Evolution Are Related to Higher Intellectual Abilities. Cereb Cortex. 25:26–34.

Garavan H, Bartsch H, Conway K, Decastro A, Goldstein RZ, Heeringa S, Jernigan T, Potter A, Thompson W, Zahs D. 2018. Recruiting the ABCD sample: Design considerations and procedures. Dev Cogn Neurosci. 32:16–22.

Gläscher J, Rudrauf D, Colom R, Paul LK, Tranel D, Damasio H, Adolphs R. 2010. Distributed neural system for general intelligence revealed by lesion mapping. Proc Natl Acad Sci U S A. 107:4705–4709.

Gottfredson LS, Deary IJ. 2004. Intelligence Predicts Health and Longevity, but Why?, CURRENT DIRECTIONS IN PSYCHOLOGICAL SCIENCE.

Grasby KL, Jahanshad N, Painter JN, Colodro-Conde L, Bralten J, Hibar DP, Lind PA, Pizzagalli F, Ching CRK, McMahon MAB, et al. 2020. The genetic architecture of the human cerebral cortex. Science (80-). 367.

Hagler DJ, Hatton S, Cornejo MD, Makowski C, Fair DA, Dick AS, Sutherland MT, Casey BJ, Barch DM, Harms MP, et al. 2019. Image processing and analysis methods for the Adolescent Brain Cognitive Development Study. Neuroimage. 116091.

Hampshire A, Highfield RR, Parkin BL, Owen AM. 2012. Fractionating Human Intelligence. Neuron. 76:1225–1237.

Harden KP, Turkheimer E, Loehlin JC. 2007. Genotype by environment interaction in adolescents’ cognitive aptitude. Behav Genet. 37:273–283.

Heaton RK, Akshoomoff N, Tulsky D, Mungas D, Weintraub S, Dikmen S, Beaumont J, Casaletto KB, Conway K, Slotkin J, et al. 2014. Reliability and validity of composite scores from the NIH Toolbox Cognition Battery in adults. J Int Neuropsychol Soc. 20:588–598.

Hickok G, Poeppel D. 2007. The cortical organization of speech processing. Nat Rev Neurosci. 8:393–402.

Hodes RJ, Insel TR, Landis SC, NIH Blueprint for Neuroscience Research. 2013. The NIH Toolbox: Setting a standard for biomedical research. Neurology. 80:S1–S1.

Horn JL, Cattell RB. 1966. Refinement and test of the theory of fluid and crystallized general intelligences. J Educ Psychol. 57:253–270.

Jernigan TL, Baaré WFC, Stiles J, Madsen KS. 2011. Postnatal brain development. Structural imaging of dynamic neurodevelopmental processes. In: Progress in Brain Research. Elsevier B.V. p. 77–92.

Jovicich J, Czanner S, Greve D, Haley E, van der Kouwe A, Gollub R, Kennedy D, Schmitt F, Brown G, MacFall J, et al. 2006. Reliability in multi-site structural MRI studies: Effects of gradient non-linearity correction on phantom and human data. Neuroimage. 30:436–443.

Kan K-J, Wicherts JM, Dolan C V., van der Maas HLJ. 2013. On the Nature and Nurture of Intelligence and Specific Cognitive Abilities. Psychol Sci. 24:2420–2428.

Lett TA, Vogel BO, Ripke S, Wackerhagen C, Erk S, Awasthi S, Trubetskoy V, Brandl EJ, Mohnke S, Veer IM, et al. 2020. Cortical Surfaces Mediate the Relationship Between Polygenic Scores for Intelligence and General Intelligence. Cereb Cortex. 30:2708–2719.

Loughnan RJ, Palmer CE, Thompson WK, Dale AM, Jernigan TL, Fan CC. 2019. Polygenic Score of Intelligence is More Predictive of Crystallized than Fluid Performance Among Children. bioRxiv. 637512.

Martin A, Schurz M, Kronbichler M, Richlan F. 2015. Reading in the brain of children and adults: a meta-analysis of 40 functional magnetic resonance imaging studies. Hum Brain Mapp. 36:1963–1981.

McDaniel MA. 2005. Big-brained people are smarter: A meta-analysis of the relationship between in vivo brain volume and intelligence. Intelligence. 33:337–346.

Menon V, Uddin LQ. 2010. Saliency, switching, attention and control: a network model of insula function. Brain Struct Funct. 214:655–667.

Mitchell BL, Cuéllar-Partida G, Grasby KL, Campos AI, Strike LT, Hwang LD, Okbay A, Thompson PM, Medland SE, Martin NG, et al. 2020. Educational attainment polygenic scores are associated with cortical total surface area and regions important for language and memory. Neuroimage. 212:116691.

Newman E, Jernigan TL, Lisdahl KM, Tamm L, Tapert SF, Potkin SG, Mathalon D, Molina B, Bjork J, Castellanos FX, et al. 2016. Go/No Go task performance predicts cortical thickness in the caudal inferior frontal gyrus in young adults with and without ADHD. Brain Imaging Behav. 10:880–892.

Newman E, Thompson WK, Bartsch H, Hagler DJ, Chen C-H, Brown TT, Kuperman JM, McCabe C, Chung Y, Libiger O, et al. 2016. Anxiety is related to indices of cortical maturation in typically developing children and adolescents. Brain Struct Funct. 221:3013–3025.

O’Leary DDM, Chou SJ, Sahara S. 2007. Area patterning of the mammalian cortex. Neuron.

Panizzon MS, Fennema-Notestine C, Eyler LT, Jernigan TL, Prom-Wormley E, Neale M, Jacobson K, Lyons MJ, Grant MD, Franz CE, et al. 2009. Distinct genetic influences on cortical surface area and cortical thickness. Cereb Cortex. 19:2728–2735.

Panizzon MS, Vuoksimaa E, Spoon KM, Jacobson KC, Lyons MJ, Franz CE, Xian H, Vasilopoulos T, Kremen WS. 2014. Genetic and Environmental Influences of General Cognitive Ability: Is g a valid latent construct? Intelligence. 43:65.

Plomin R, Spinath FM. 2004. Intelligence: Genetics, Genes, and Genomics. J Pers Soc Psychol. 86:112–129.

Posthuma D, Baaré WFC, Hulshoff Pol HE, Kahn RS, Boomsma DI, De Geus EJC. 2003. Genetic Correlations Between Brain Volumes and the WAIS-III Dimensions of Verbal Comprehension, Working Memory, Perceptual Organization, and Processing Speed. Twin Res. 6:131–139.

Rakic P, Ayoub AE, Breunig JJ, Dominguez MH. 2009. Decision by division: making cortical maps. Trends Neurosci.

Reardon PK, Seidlitz J, Vandekar S, Liu S, Patel R, Park MTM, Alexander-Bloch A, Clasen LS, Blumenthal JD, Lalonde FM, et al. 2018. Normative brain size variation and brain shape diversity in humans. Science (80-). 360:1222–1227.

Reddan MC, Lindquist MA, Wager TD. 2017. Effect size estimation in neuroimaging. JAMA Psychiatry.

Savage JE, Jansen PR, Stringer S, Watanabe K, Bryois J, De Leeuw CA, Nagel M, Awasthi S, Barr PB, Coleman JRI, et al. 2018. Genome-wide association meta-analysis in 269,867 individuals identifies new genetic and functional links to intelligence. Nat Genet. 50:912–919.

Schmitt JE, Neale MC, Clasen LS, Liu S, Seidlitz J, Pritikin JN, Chu A, Wallace GL, Lee NR, Giedd JN. 2019. A comprehensive quantitative genetic analysis of cerebral surface area in youth. J Neurosci. 39:3028–3040.

Schnack HG, van Haren NEM, Brouwer RM, Evans A, Durston S, Boomsma DI, Kahn RS, Hulshoff Pol HE. 2015. Changes in Thickness and Surface Area of the Human Cortex and Their Relationship with Intelligence. Cereb Cortex. 25:1608–1617.

Shadrin AA, Kaufmann T, van der Meer D, Palmer CE, Makowski C, Loughnan RM, Jernigan TL, Seibert TM, Hagler DJ, Smeland OB, et al. 2020. Multivariate genome-wide association study identifies 1735 unique genetic loci associated with cortical morphology. bioRxiv. 2020.10.22.350298.

Shaw P, Greenstein D, Lerch J, Clasen L, Lenroot R, Gogtay N, Evans A, Rapoport J, Giedd J. 2006. Intellectual ability and cortical development in children and adolescents. Nature. 440:676–679.

Sowell ER, Thompson PM, Leonard CM, Welcome SE, Kan E, Toga AW. 2004. Longitudinal Mapping of Cortical Thickness and Brain Growth in Normal Children. J Neurosci. 24:8223–8231.

Spearman C. 1904. “General Intelligence,” Objectively Determined and Measured. Am J Psychol. 15:201.

Taylor KS, Seminowicz DA, Davis KD. 2009. Two systems of resting state connectivity between the insula and cingulate cortex. Hum Brain Mapp. 30:2731–2745.

Van Der Maas H, Kan K-J, Marsman M, Stevenson CE. 2017. Network Models for Cognitive Development and Intelligence. J Intell. 5:16.

Van Der Maas HLJ, Dolan C V, Grasman RPPP, Wicherts JM, Huizenga HM, Raijmakers MEJ. 2006. A Dynamical Model of General Intelligence: The Positive Manifold of Intelligence by Mutualism.

van der Meer D, Frei O, Kaufmann T, Shadrin AA, Devor A, Smeland OB, Thompson WK, Fan CC, Holland D, Westlye LT, et al. 2020. Understanding the genetic determinants of the brain with MOSTest. Nat Commun. 11:1–9.

Vuoksimaa E, Panizzon MS, Chen C-H, Fiecas M, Eyler LT, Fennema-Notestine C, Hagler DJ, Fischl B, Franz CE, Jak A, et al. 2015. The Genetic Association Between Neocortical Volume and General Cognitive Ability Is Driven by Global Surface Area Rather Than Thickness. Cereb Cortex. 25:2127–2137.

Vuoksimaa E, Panizzon MS, Chen C-H, Fiecas M, Eyler LT, Fennema-Notestine C, Hagler DJ, Franz CE, Jak AJ, Lyons MJ, et al. 2016. Is bigger always better? The importance of cortical configuration with respect to cognitive ability. Neuroimage. 129:356–366.

Walhovd KB, Krogsrud SK, Amlien IK, Bartsch H, Bjørnerud A, Due-Tønnessen P, Grydeland H, Hagler DJ, Håberg AK, Kremen WS, et al. 2016. Neurodevelopmental origins of lifespan changes in brain and cognition. Proc Natl Acad Sci U S A. 113:9357–9362.

Wechsler D. 1946. The measurement of adult intelligence (3rd ed.). Baltimore: Williams & Wilkins Co.

Winkler AM, Kochunov P, Blangero J, Almasy L, Zilles K, Fox PT, Duggirala R, Glahn DC. 2010. Cortical thickness or grey matter volume? The importance of selecting the phenotype for imaging genetics studies. Neuroimage. 53:1135–1146.

Winkler AM, Webster MA, Vidaurre D, Nichols TE, Smith SM. 2015. Multi-level block permutation. Neuroimage. 123:253–268.

Zhao W, Palmer CE, Thompson WK, Chaarani B, Garavan HP, Casey BJ, Jernigan TL, Dale AM, Fan CC. 2020. Individual Differences in Cognitive Performance Are Better Predicted by Global Rather Than Localized BOLD Activity Patterns Across the Cortex. Cereb Cortex.

