## Supplementary Material for "Distinct regionalization patterns of cortical morphology are associated with cognitive performance across different domains"


- Extended Methods ........................................................................................... 2-5
- Supplementary Tables 1-3 ............................................................................... 6-8
- Supplementary Figures 1-5 ............................................................................. 9-13

### Extended Methods

**Sample.** There was a large number of subjects with missing income data (n=1018), therefore missing values were imputed by taking the median income value across participants from the same testing site. The sources of other missing data were as follows: incomplete across all demographic variables (n=189), incomplete across all cognitive measures (n=944), unavailable T1-weighted MRI scan for reasons outlined in the ABCD release notes (e.g., did not get scanned, motion artefacts) (n=339) and imaging data that was made available but did not pass the free-surfer QC flag (n=462). These missing data values are not mutually exclusive. The permutation testing was dependent on having multiple families with the same number of children, therefore the single family with 5 children was excluded from these analyses. Supplementary Table 1 displays the names of each variable used in these analyses from data release 2.0.1. Supplementary Table 2 shows the demographic characteristics of the sample as a function of cognitive performance.

**Neurocognitive battery**, Below are details of each of the cognitive tasks analyzed in this study.

***Toolbox Oral Reading Recognition Task® (TORRT):*** measured language decoding and reading. Children were asked to read aloud single letters or words presented in the center of an iPad screen. The research assistant marked pronunciations as correct or incorrect. Extensive training was given prior to administering the test battery. Item difficulty was modulated using computerized adaptive testing (CAT).

***Toolbox Picture Vocabulary Task® (TPVT)***: a variant of the Peabody Picture Vocabulary Test (PPTV), measured language and vocabulary comprehension. Four pictures were presented on an iPad screen as a word was played through the iPad speaker. The child was instructed to point to the picture, which represented the concept, idea or object name heard. CAT was implemented to control for item difficulty and avoid floor or ceiling effects.

***Toolbox Pattern Comparison Processing Speed Test® (TPCPST)***: measured processing speed. Children were shown two images and asked to determine if they were identical or different by touching the appropriate response button on the screen. This test score is the sum of the number of items completed correctly in the time given.

***Toolbox List Sorting Working Memory Test® (TLSWMT)***: measured working memory. Children heard a list of words alongside pictures of each word and were instructed to repeat the list back in order of their actual size from smallest to largest. The list started with only 2 items and a single category (e.g., animals). The number of items increased with each correct answer to a maximum of seven. The child then progressed to the next stage in which two different categories were interleaved. At this stage children were required to report the items back in size order from the first category followed by the second category. Children were always given two opportunities to repeat the list correctly before the experimenter scored the trial as incorrect.

***Toolbox Picture Sequence Memory Test® (TPSMT)***: measured episodic memory. On each trial, children were shown a series of fifteen pictures in a particular sequence. The pictures illustrated activities or events within a particular setting (e.g., going to the park), and as each appeared on the screen a pre-recorded narrative briefly described the content of the picture. Participants were instructed to arrange the pictures in the original sequence in which they were shown. The Rey-Auditory Verbal Learning Task was also included in the ABCD neurocognition battery as a more comprehensive measure of episodic memory.

***Toolbox Flanker Task® (TFT):*** measured executive function, attentional and inhibitory control. This adaptation of the Eriksen Flanker task(Eriksen and Eriksen 1974) captures how readily a participant is influenced by the congruency of stimuli surrounding a target. On each trial a target arrow was presented in the center of the iPad screen facing to the left or right and was flanked by two additional arrows on both sides. The surrounding arrows were either facing in the same (congruent) or different (incongruent) direction to the central target arrow. The participant was instructed to push a response button to indicate the direction of the central target arrow. Accuracy and reaction time scores were combined to produce a total score of executive attention, such that higher scores indicate a greater ability to attend to relevant information and inhibit incorrect responses.

***Toolbox Dimensional Change Card Sort Task® (TDCCS)***: measured executive function and cognitive flexibility. On each trial, the participant was presented with two objects at the bottom of the iPad screen and a third object in the middle. The participant was asked to sort the third object by matching it to one of the bottom two objects based on either color or shape. In the first block participants matched based on one dimension and in the second block they switched to the other dimension. In the final block, the sorting dimension alternated between trials pseudorandomly. The total score was calculated based on speed and accuracy.

***Rey-Auditory Verbal Learning Task (RAVLT)***: measures auditory learning, recall and recognition. Participants listened to a list of 15 unrelated words and were asked to immediately recall these after each of five learning trials. A second unrelated list was then presented, and participants were asked to recall as many words as possible from the second list and then recall words again from the initial list. Following a delay of 30 minutes (during which other non-verbal tasks from the cognitive battery are administered), longer-term retention was measured using recall and recognition. This task was administered via an iPad using the Q-interactive platform of Pearson assessments (Daniel et al. 2014). In the current study, the total number of items correctly recalled across the five learning trials was summed to produce a measure of auditory verbal learning.

***Little Man Task (LMT):*** measures visuospatial processing involving mental rotation with varying degrees of difficulty (Acker 1982). A rudimentary male figure holding a briefcase in one hand was presented on an iPad screen. The figure could appear in one of four positions: right side up vs upside down and either facing the participant or with his back to the participant. The briefcase could be in either hand. Participants indicated which hand the briefcase was in using one of two buttons. Performance across the 32 trials was measured by the percentage of trials in which the child responded correctly. This was divided by the average reaction time to complete the task (in seconds) to produce a measure of efficiency of visuospatial processing. This was the dependent variable analyzed in this study.

***WISC-V Matrix reasoning***. Nonverbal reasoning was measured using an automated version of the Matrix Reasoning subtest from the Weschler Intelligence Test for Children-V (WISC-V; Weschler, 2014). On each trial the participant was presented with a series of visuospatial stimuli, which was incomplete. The participant was instructed to select the next stimulus in the sequence from four alternatives. There were 32 possible trials and testing ended when the participant failed three consecutive trials. The total raw score, used in the current study, was the total number of trials completed correctly.

**MRI acquisition and image pre-processing.** The T1w acquisition (1 mm isotropic) was a 3D T1w inversion prepared RF-spoiled gradient echo scan using prospective motion correction, when available (White et al. 2010; Tisdall et al. 2012) (echo time = 2.88 ms, repetition time = 2500 ms, inversion time = 1060 ms, flip angle = 8°, FOV = 256x256, FOV phase = 100%, slices = 176). Pre-processing of all MRI data for ABCD was conducted using in-house software at the Center for Multimodal Imaging and Genetics (CMIG) at University of California San Diego (UCSD) as outlined in Hagler et al (Hagler et al. 2019). Manual quality control was performed prior to the full image pre-processing and structural scans with poor image quality as well as those that did not pass FreeSurfer QC were excluded from all analyses. Brain segmentation and cortical surface reconstruction were completed using FreeSurfer v5.3.0 (Dale et al. 1999; Fischl et al. 1999). T1-weighted structural images were corrected for distortions caused by gradient nonlinearities, coregistered, averaged, and rigidly resampled into alignment with an atlas brain. See previous publications for details of the surface based cortical reconstruction segmentation procedures (Dale et al. 1999; Fischl et al. 1999, 2004; Fischl and Dale 2000; Jovicich et al. 2006). In brief, a 3D model of the cortical surface was constructed for each subject. This included segmentation of the white matter (WM), tessellation of the gray matter (GM)/WM boundary, inflation of the folded, tessellated surface, and correction of topological defects. Measures of cortical thickness at each vertex were calculated as the shortest distance between the reconstructed GM/WM and pial surfaces (Fischl and Dale 2000). To calculate cortical surface area, a standardized tessellation was mapped to the native space of each subject using a spherical atlas registration, which matched the cortical folding patterns across subjects. Surface area of each point in atlas space was calculated as the area of each triangle. This generated a continuous vertex-wise measure of relative areal expansion or contraction. Cortical maps were smoothed using a Gaussian kernel of 20 mm full-width half maximum (FWHM) and mapped into standardized spherical atlas space.

#### Statistical Analysis

***Vertex-wise effect size maps.*** All behavioral variables were standardized (z-scored) prior to analysis as was the vertex-wise brain data. We applied a general linear model (GLM) associating a given behavior $y$ from a set of covariates $W$ and the vertex-wise morphology data, $X_v$

$$y = \alpha_{0,v} + \boldsymbol{W}\alpha_v + X_v\beta_v + \varepsilon_v$$

Here, $W$ represents a standardized $N \ x \ m$ matrix of covariates of no interest where $N$ represents the number of subjects and $m$ the number of covariates. $X_v$ denotes the standardised $N \ x \ 1$ vector of imaging data for the $v$th vertex. This GLM was applied univariately at each vertex $v = 1, \ldots V$. Let $\boldsymbol{\beta} = (\beta_1, \ldots, \beta_V)'$ and α denote the $V \ x \ 1$ and $m \ x \ 1$ vectors of parameters of interest and no interest, respectively. For visualisation of these effects the standardized $\boldsymbol{\beta}$ coefficients were plotted across vertices. All vertexwise effect size maps were calculated with and without including race/ethnicity, income and parental education as predictors in the covariates matrix, $W$.

***Determining the significance of the effect size maps using an omnibus test.*** Assuming that $\boldsymbol{z}$ is asymptotically drawn from a multivariate normal distribution and $\widehat{\boldsymbol{R}}$ is a consistent estimator of the covariance matrix $\boldsymbol{R}$, the MOSTest statistic would have an approximate Chi-squared distribution (a special case of the gamma distribution) under the null hypothesis. However, we relax this assumption and instead compute its p-value using a hybrid permutation procedure as with the other statistics. The covariance matrix $\widehat{\boldsymbol{R}}$ was estimated from the $V \ x \ L$ matrix of permuted Wald statistics, $\boldsymbol{Z_{perm}}$, where $L$ denotes the number of permutations. This was then regularized using truncated singular value decomposition (SVD) to ensure the covariance matrix was invertible. The truncation parameter, *kmax*, was determined as the point in which the rate of decrease in magnitude of the eigenvalues decreased substantially towards 0 (quantified at the point in which all subsequent eigenvalues were below 0.1; supplementary figure 3).

For the permutation procedure, families with the same number of siblings were allowed to be shuffled as a whole and siblings within a family were allowed to be shuffled with each other. This was necessary to account for the joint distribution of the observed data due to the relatedness of the ABCD sample. Permutations that adhered to this shuffling scheme were generated using the PALM toolbox (Winkler et al. 2014). Permutations were conducted using the Freedman-Lane procedure(Freedman and Lane 1983). This permutation procedure was used to determine the distribution of each test statistic under the global null hypothesis $H_0$. We rejected $H_0$ if the observed test statistic was greater than the value of the permuted test statistic at the critical threshold corresponding to an alpha level of 0.0038 (0.05 corrected for the 13 cognitive tests analyzed). To extrapolate the p-values beyond the range that can be directly estimated from the permutations, we fit tail of the permuted null test statistics to specific analytic forms: gamma distribution for the MOSTest test statistic, and Weibull distribution for the -log(min-p). Supplementary figure 2 shows the fit of the fitted distributions to the permuted data for each test statistic. Fits were best for the MOSTest compared to the other test statistics, which suggests that extrapolated p-values for the other associations may be inflated.

***Quantifying the magnitude of the association between brain structure and cognition using a polyvertex score (PVS).*** The association between each imaging phenotype and each cognitive task was modelled using the mass univariate approach such that the behavior of interest was predicted independently at each vertex using a general linear model (GLM). The PVS predicting behavior from cortical morphometry was then computed for each subject as the product sum of the estimated effects and the pre-residualized cortical morphometry vector. This measure thus harnesses the explanatory power of all of the vertices with respect to behavior. The computed PVS was then compared with the observed behavior in order to provide an estimate for how much variance in the observed behavior can be predicted using the vertex-wise imaging phenotype. In order to generate an unbiased PVS for every subject we used a leave-one-out 10-fold cross-validation procedure. The cortical mass univariate effects were estimated in the training set (90% of full sample) and multiplied with the imaging phenotype of participants in the test set (10% of full sample). This was repeated 10 times for each fold until a PVS was calculated for every participant in the full sample. The subjects in each fold were randomly selected based on unique family IDs, such that subjects within the same family were always within the same fold. The association between the imaging phenotype and behavior across the whole sample was calculated as the squared correlation ($R^2$) between the observed behavior and the predicted behavior (the PVS). This process was repeated for four imaging phenotypes: CSA, relative CSA (controlling for total CSA), CTH. relative CTH (controlling for mean CTH).

In order to explore the proportion of shared variability in cognitive performance explained by brain structure and the demographic variables, we generated several linear models with differing predictors. These models were generated separately for each imaging modality (when these were included) and with either the fluid or crystallized scores as the dependent variable. Each model was trained on 90% of the sample and tested in a 10% hold out set within a 10-fold cross validation framework to produce a robust, out-of-sample $R^2$. The models are outlined in figure 7. Where indicated both the dependent and independent variables were pre-residualized for the stated variables (in parentheses), in order to remove the variance associated with those variables and calculate the unique variance associated only with the variables of interest. This method was used to partition the shared and unique variance across the measures of interest. It is important to note, this method offers an approximation of the unique behavioral variance associated with these variables of interest, because these variables are not completely orthogonal to one another. Here we always pre-residualize for age, sex and scanner first as nuisance variables except for in the full model. The full model (model 1) gives an estimate of the maximal variance explained when all of the variables measured are included in a model together and thus their covariance is accounted for when estimating the model $R^2$.

**Supplementary References**


Acker W. 1982. A computerized approach to psychological screening—The Bexley-Maudsley Automated Psychological Screening and The Bexley-Maudsley Category Sorting Test, International Journal of Human-computer Studies / International Journal of Man-machine Studies - IJMMS.

Dale AM, Fischl B, Sereno MI. 1999. Cortical Surface-Based Analysis. Neuroimage. 9:179–194.

Daniel MH, Wahlstrom D, Zhang O. 2014. Equivalence of Q-interactive™ and Paper Administrations of Cognitive Tasks: WISC ®-V Q-interactive Technical Report 8.

Eriksen BA, Eriksen CW. 1974. Effects of noise letters upon the identification of a target letter in a nonsearch task. Percept Psychophys. 16:143–149.

Fischl B, Dale AM. 2000. Measuring the thickness of the human cerebral cortex from magnetic resonance images. Proc Natl Acad Sci U S A. 97:11050–11055.

Fischl B, Salat DH, van der Kouwe AJW, Makris N, Ségonne F, Quinn BT, Dale AM. 2004. Sequence-independent segmentation of magnetic resonance images. Neuroimage. 23:S69--S84.

Fischl B, Sereno MI, Dale AM. 1999. Cortical Surface-Based Analysis II: Inflation, Flattening, and a Surface-Based Coordinate System.

Freedman D, Lane D. 1983. A Nonstochastic Interpretation of Reported Significance Levels. J Bus Econ Stat. 1:292.

Hagler DJ, Hatton S, Cornejo MD, Makowski C, Fair DA, Dick AS, Sutherland MT, Casey BJ, Barch DM, Harms MP, et al. 2019. Image processing and analysis methods for the Adolescent Brain Cognitive Development Study. Neuroimage. 116091.

Jovicich J, Czanner S, Greve D, Haley E, van der Kouwe A, Gollub R, Kennedy D, Schmitt F, Brown G, MacFall J, et al. 2006. Reliability in multi-site structural MRI studies: Effects of gradient non-linearity correction on phantom and human data. Neuroimage. 30:436–443.

Tisdall MD, Hess AT, Reuter M, Meintjes EM, Fischl B, van der Kouwe AJW. 2012. Volumetric navigators for prospective motion correction and selective reacquisition in neuroanatomical MRI. Magn Reson Med. 68:389–399.

Weschler D. 2014. Weschler Intelligence Scale for Children, 5th ed. Pearson, Bloomington, MN.

White N, Roddey C, Shankaranarayanan A, Han E, Rettmann D, Santos J, Kuperman J, Dale A. 2010. PROMO: Real-time prospective motion correction in MRI using image-based tracking. Magn Reson Med. 63:91–105.

Winkler AM, Ridgway GR, Webster MA, Smith SM, Nichols TE. 2014. Permutation inference for the general linear model. Neuroimage. 92:381–397.

| Type of Variable | Variable | NDA/DEAP Variable Name | Instrument |
|---|---|---|---|
| ***Covariates of no interest*** | Age | interview_age | Developmental history questionnaire (dhx01) |
| | Sex | sex | Developmental history questionnaire (dhx01) |
| | Self-declared race/ethnicity | race.4level**, hisp** | Parent Demographics Survey (pdem02) – computed DEAP variable |
| | Income | household.income | Parent Demographics Survey (pdem02) – computed DEAP variable |
| | Parental education | high.educ | Parent Demographics Survey (pdem02) – computed DEAP variable |
| | Scanner ID | mri_info_deviceserialnumber | MRI Info (abcd_mri01) |
| ***Family ID*** | Family | rel_family_id | ACS Post Stratification Weights (acspsw02) |
| ***MRI QC*** | Freesurfer QC | fsqc_qc | FreeSurfer QC (freesqc01) |
| ***Cognitive Measures*** | Toolbox Oral Reading Recognition | nihtbx_reading_uncorrected | Youth NIH TB Summary Scores (abcd_tbss01) |
| | Toolbox Picture Vocabulary Task | nihtbx_picvocab_uncorrected | Youth NIH TB Summary Scores (abcd_tbss01) |
| | Toolbox Flanker Task | nihtbx_flanker_uncorrected | Youth NIH TB Summary Scores (abcd_tbss01) |
| | Toolbox Dimensional Card Sorting Task | nihtbx_cardsort_uncorrected | Youth NIH TB Summary Scores (abcd_tbss01) |
| | Toolbox Pattern Speed Task | nihtbx_pattern_uncorrected | Youth NIH TB Summary Scores (abcd_tbss01) |
| | Toolbox Picture Sequence Task | nihtbx_picture_uncorrected | Youth NIH TB Summary Scores (abcd_tbss01) |
| | Toolbox List Working Memory Task | nihtbx_list_uncorrected | Youth NIH TB Summary Scores (abcd_tbss01) |
| | Toolbox Fluid Composite Score | nihtbx_fluidcomp_uncorrected | Youth NIH TB Summary Scores (abcd_tbss01) |
| | Toolbox Crystallised Composite Score | nihtbx_cryst_uncorrected | Youth NIH TB Summary Scores (abcd_tbss01) |
| | Matrix reasoning | pea_wiscv_trs | Pearson Scores (abcd_ps01) |
| | Little Man Task | lmt_scr_efficiency | Little Man Task Summary Scores (lmtp201) |
| | RAVLT | pea_ravlt_sd_trial_i_tc+pea_ravlt_sd_trial_ii_tc+pea_ravlt_sd_trial_iii_tc+pea_ravlt_sd_trial_iv_tc+pea_ravlt_sd_trial_v_tc | Pearson Scores (abcd_ps01) |

**Supplementary Table 1.** List of all variables used in this study with corresponding variable names and the instruments in which they can be found in the ABCD data dictionary and data release. Only participants with complete data on all of these variables, available T1-weighted MRI scans and whose imaging data passed the freesurfer QC flag (fsqc_qc=1) were included in the analyses.** *We computed race/ethnicity by changing the race of all those who endorsed being Hispanic to 'Hispanic' to create a 5-level race/ethnicity variable as seen in release 1.1 and 2.0.*

| | |
|---|---|
| ***n*** | *10145* |
| **Toolbox Fluid Composite Score (mean (SD))** | 91.76 (10.54) |
| **Toolbox Crystallised Composite Score (mean (SD))** | 86.50 (7.03) |
| ***Age (mean (sd))*** | 119.02 (7.48) |
| ***Sex = M (%)*** | 5288 (52.1) |
| ***Parent education (%)*** | |
| ***< HS Diploma*** | 460 ( 4.5) |
| ***HS Diploma/GED*** | 942 ( 9.3) |
| ***Some College*** | 2624 (25.9) |
| ***Bachelor*** | 2611 (25.7) |
| ***Post Graduate Degree*** | 3508 (34.6) |
| ***Income (%)*** | |
| ***<=50k*** | 2676 (28.7) |
| ***>50k<=100k*** | 2661 (28.6) |
| ***>100k*** | 3971 (42.7) |
| ***Race/ethnicity (%)*** | |
| ***White*** | 5411 (53.3) |
| ***Black*** | 1484 (14.6) |
| ***Asian*** | 211 ( 2.1) |
| ***Other/Mixed*** | 989 ( 9.7) |
| ***Hispanic*** | 2050 (20.2) |
| ***Scanner ID (%)*** | |
| ***HASH03db707f*** | 423 ( 4.2) |
| ***HASH11ad4ed5*** | 480 ( 4.7) |
| ***HASH1314a204*** | 506 ( 5.0) |
| ***HASH311170b9*** | 339 ( 3.3) |
| ***HASH31ce566d*** | 43 ( 0.4) |
| ***HASH3935c89e*** | 959 ( 9.5) |
| ***HASH48f7cbc3*** | 29 ( 0.3) |
| ***HASH4b0b8b05*** | 387 ( 3.8) |
| ***HASH4d1ed7b1*** | 352 ( 3.5) |
| ***HASH5ac2b20b*** | 462 ( 4.6) |
| ***HASH5b0cf1bb*** | 559 ( 5.5) |
| ***HASH5b2fcf80*** | 255 ( 2.5) |
| ***HASH65b39280*** | 307 ( 3.0) |
| ***HASH69f406fa*** | 130 ( 1.3) |
| ***HASH6b4422a7*** | 318 ( 3.1) |
| ***HASH7911780b*** | 358 ( 3.5) |
| ***HASH7f91147d*** | 86 ( 0.8) |
| ***HASH96a0c182*** | 535 ( 5.3) |
| ***HASHa3e45734*** | 291 ( 2.9) |
| ***HASHb640a1b8*** | 471 ( 4.6) |
| ***HASHc3bf3d9c*** | 439 ( 4.3) |
| ***HASHc9398971*** | 199 ( 2.0) |
| ***HASHd422be27*** | 419 ( 4.1) |
| ***HASHd7cb4c6d*** | 497 ( 4.9) |
| ***HASHdb2589d4*** | 495 ( 4.9) |
| ***HASHe3ce02d3*** | 86 ( 0.8) |
| ***HASHe4f6957a*** | 505 ( 5.0) |
| ***HASHe76e6d72*** | 15 ( 0.1) |
| ***HASHfeb7e81a*** | 200 ( 2.0) |

**Supplementary Table 2. Descriptive characteristics of the analyzed sample.**

| *Full sample partial r* | | | | | | | | | |
|---|---|---|---|---|---|---|---|---|---|
| | | Not controlling for Socio-demographics | | Controlling for Socio-demographics | | Not controlling for Socio-demographics | | Controlling for Socio-demographics | |
| **DV** | **IV** | **CSA $PVS_U$** | **TOTAL CSA** | **CSA $PVS_U$** | **TOTAL CSA** | **CTH $PVS_U$** | **MEAN CTH** | **CTH $PVS_U$** | **MEAN CTH** |
| **Fluid** | **PC1_PVS** | 0.13 | 0.17 | 0.06 | 0.08 | 0.16 | 0.02 | 0.10 | -0.02 |
| **Fluid** | **F_PVS** | 0.12 | 0.17 | 0.07 | 0.08 | 0.16 | 0.02 | 0.10 | -0.02 |
| **Fluid** | **C_PVS** | 0.09 | 0.17 | 0.02 | 0.08 | 0.14 | 0.02 | 0.07 | -0.02 |
| **Cryst** | **PC1_PVS** | 0.14 | 0.27 | 0.05 | 0.15 | 0.19 | 0.06 | 0.08 | -0.01 |
| **Cryst** | **F_PVS** | 0.11 | 0.27 | 0.02 | 0.15 | 0.19 | 0.06 | 0.07 | -0.01 |
| **Cryst** | **C_PVS** | 0.14 | 0.27 | 0.06 | 0.15 | 0.19 | 0.06 | 0.09 | -0.01 |
| **PC1** | **PC1_PVS** | 0.17 | 0.25 | 0.07 | 0.13 | 0.21 | 0.05 | 0.12 | -0.02 |
| **PC1** | **F_PVS** | 0.15 | 0.25 | 0.06 | 0.13 | 0.22 | 0.05 | 0.11 | -0.02 |
| **PC1** | **C_PVS** | 0.14 | 0.25 | 0.05 | 0.13 | 0.20 | 0.05 | 0.09 | -0.02 |
| ***Cross-validated %$R^2$*** | | | | | | | | | |
| **Fluid** | **PC1_PVS** | 1.60 | 2.80 | 0.31 | 0.54 | 2.38 | 0.04 | 1.02 | 0.02 |
| **Fluid** | **F_PVS** | 1.41 | 2.82 | 0.39 | 0.55 | 2.53 | 0.03 | 0.89 | 0.03 |
| **Fluid** | **C_PVS** | 0.83 | 2.79 | 0.00 | 0.53 | 1.95 | 0.03 | 0.47 | 0.02 |
| **Cryst** | **PC1_PVS** | 1.97 | 7.26 | 0.20 | 2.19 | 3.68 | 0.37 | 0.57 | 0.02 |
| **Cryst** | **F_PVS** | 1.13 | 7.19 | 0.01 | 2.21 | 3.56 | 0.37 | 0.44 | 0.07 |
| **Cryst** | **C_PVS** | 1.95 | 7.29 | 0.32 | 2.20 | 3.51 | 0.36 | 0.73 | 0.03 |
| **PC1** | **PC1_PVS** | 2.88 | 6.35 | 0.52 | 1.65 | 4.50 | 0.20 | 1.31 | 0.00 |
| **PC1** | **F_PVS** | 2.17 | 6.32 | 0.37 | 1.64 | 4.63 | 0.17 | 1.10 | 0.00 |
| **PC1** | **C_PVS** | 2.01 | 6.29 | 0.21 | 1.63 | 3.91 | 0.22 | 0.82 | 0.01 |
| **PC1** | **F_PVS+C_PVS** | 2.87 | 6.34 | 0.50 | 1.64 | 4.67 | 0.20 | 1.28 | 0.02 |

**Supplementary Table 3. Unique associations between behavior and regional and global structural phenotypes.** Associations are shown as both partial correlation coefficients in the full sample (top) and cross-validated, out-of-sample %$R^2$ (bottom) after residualizing for covariates of no interest only (age, sex, scanner ID) and additionally residualizing for sociodemographic variables (household income, race/ethnicity, parental education). Each row shows a different model. PC1, fluid and crystallized composite scores were predicted by either relative CSA/CTH (the PC1, fluid or crystallized $PVS_U$) and in a separate model total CSA/mean CTH residualized for the respective $PVS_U$ indicated in column 'IV'. Associations for relative CSA and Total CSA, and relative CTH and mean CTH, are therefore unique. The $F\_PVS_U$ and $C\_PVS_U$ were less predictive of the behavior they were not trained on highlighting the differences in the regionalization associations. They predicted a similar proportion of variance in PC1. The final row shows a model in which PC1 was predicted by both $F\_PVS_U$ and $C\_PVS_U$ in the same model. When controlling for sociodemographic factors, the $R^2$ was very similar to the sum of the $R^2$ of each PVS predicting PC1 independently, which suggests these imaging measures are predicting non-overlapping variance in PC1. This was more evident for CSA than CTH due to the increased similarity in the CTH regionalization association patterns. When not controlling for sociodemographic factors, the $F\_PVS_U$ and $C\_PVS_U$ predicted overlapping variance in PC1 indicative of the shared variance across the cognitive, brain and sociodemographic variables.

A

| Cognitive Measures | PC1 Loadings |
|---|---|
| nihtbx_picvocab_uncorrected | 0.67 |
| nihtbx_flanker_uncorrected | 0.58 |
| nihtbx_list_uncorrected | 0.68 |
| nihtbx_cardsort_uncorrected | 0.65 |
| nihtbx_pattern_uncorrected | 0.50 |
| nihtbx_picture_uncorrected | 0.56 |
| nihtbx_reading_uncorrected | 0.67 |
| pea_ravlt_ld | 0.65 |
| lmt_scr_efficiency | 0.44 |
| pea_wiscv_trs | 0.64 |
| **Proportion Variance** | **0.37** |

B

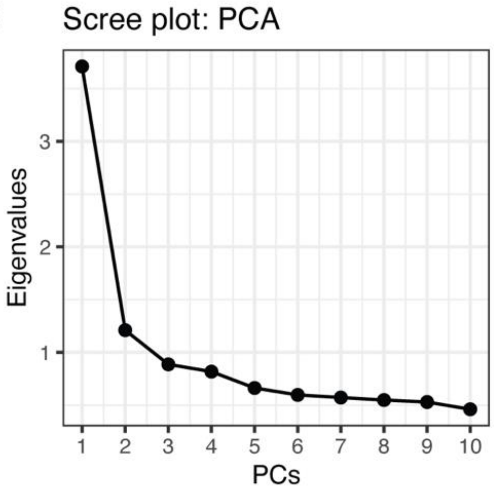


**Supplementary Figure 1.** A) Loadings for each of the cognitive tasks and the first unrotated principal component (PC1) from the PCA. B) Scree plot showing the eigenvalues for the top 10 components of the PCA. PC1 explained 37% of the variance across tasks.

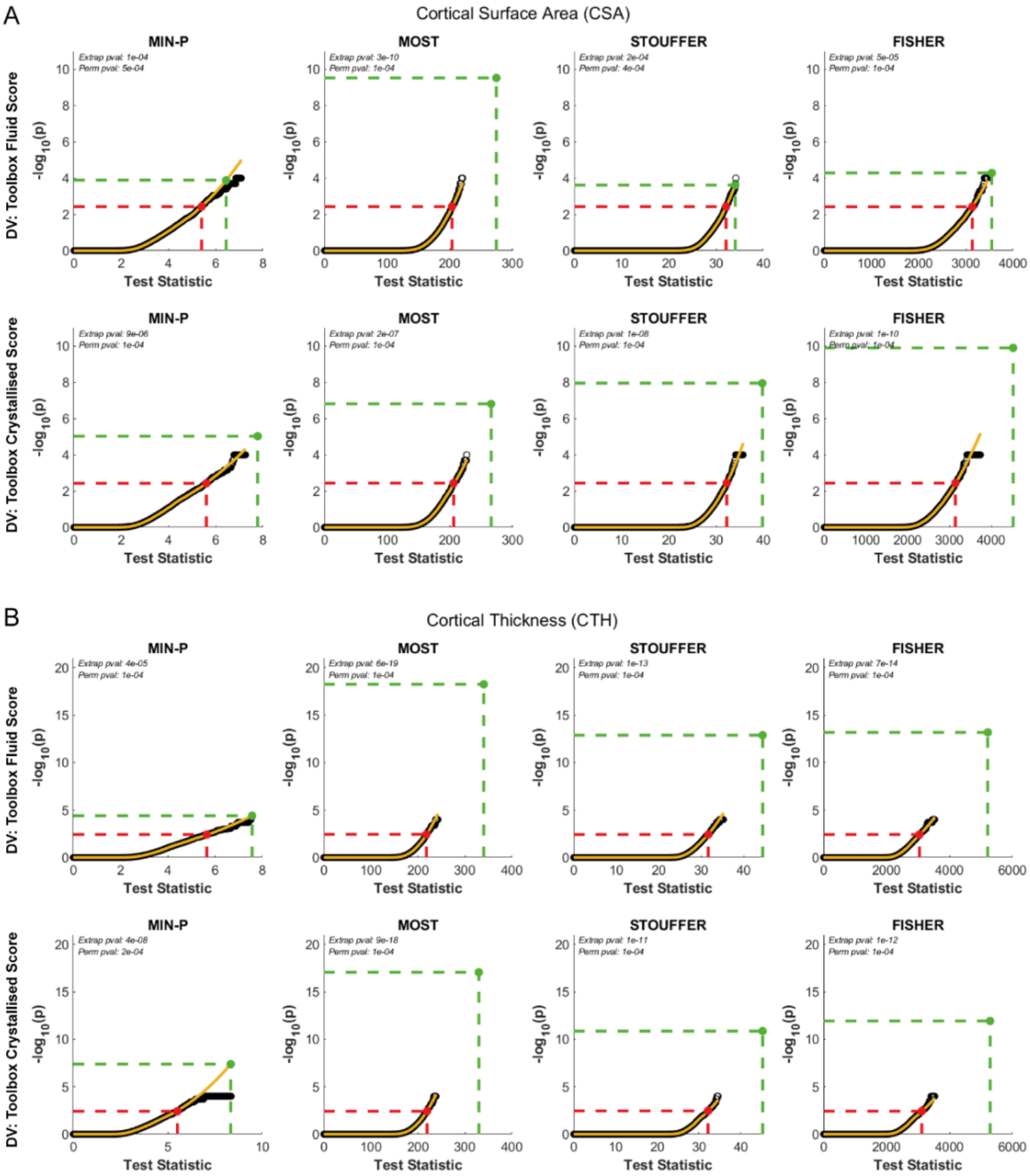


**Supplementary Figure 2. Cumulative density functions (CDF) for permuted and fitted data across all associations.** The CDF of the permuted test statistic (black) and the predicted CDF (yellow) after fitting either a Weibull distribution (column 1) to the permuted univariate statistics or gamma distribution to the permuted multivariate statistics (columns 2-4). The fitted CDF was extrapolated to provide an approximated p-value for the observed statistic. For each association and across each of the test statistics, the observed statistics for the associations between cortical morphology and cognition were statistically significant i.e., the observed statistic (green) was further in the tail of the null distribution (black) than the FWE-corrected threshold (red). The magnitude of effects were larger for the multivariate statistics than the univariate case likely due to the distributed nature of the brain-behavior effects across the cortex.

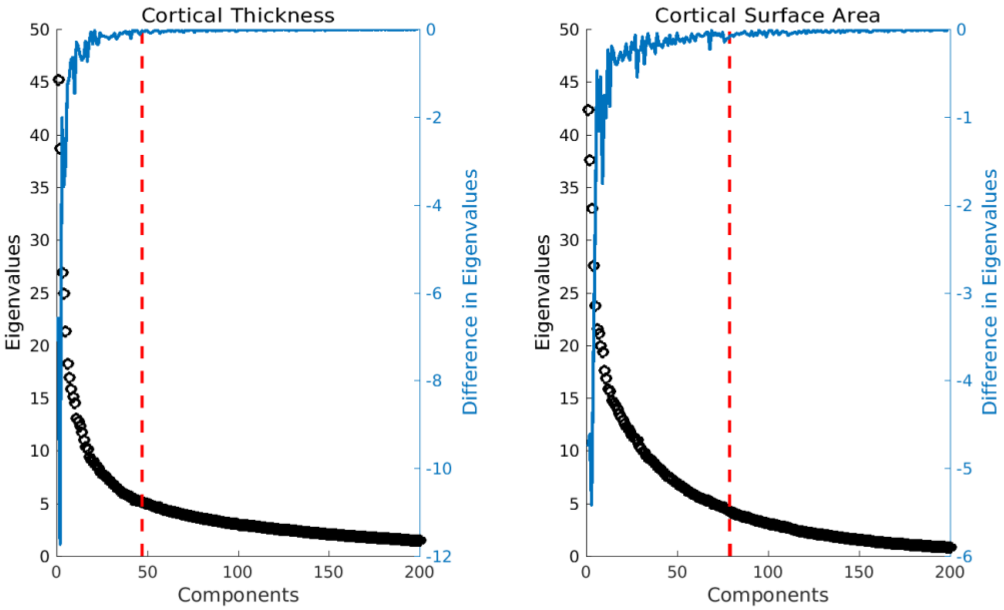


**Supplementary Figure 3.** Eigenvalues (black) for 200 of the 1284 components from the SVD of the covariance matrix for cortical thickness (left) and cortical surface area (right). The vertexwise covariance matrix was estimated based on the z statistics from 10,000 permuted vertexwise associations between cortical morphology and cognition. This used a different permutation scheme to the main results in this paper to avoid any biases. The vertexwise covariance matrix was used for the calculation of the MOSTest test statistic. To ensure the matrix was invertible the high dimensional matrix was regularized using a truncated SVD approach based on spectral filtering of the covariance matrix using all eigenvalues above the red dotted line (truncation parameter, k). k was determined based on the point at which the rate of change in the magnitude of the eigenvalues decreased towards zero (blue).

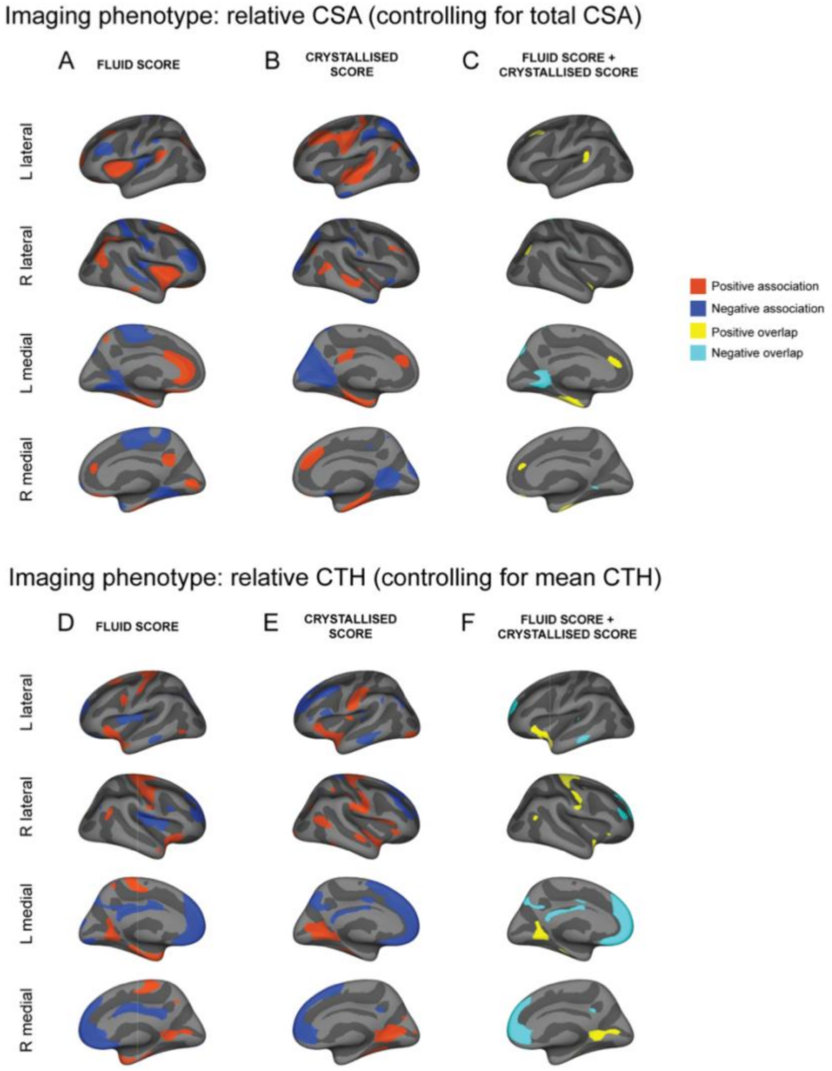


**Supplementary Figure 4. Similarity maps for the vertexwise structure-function associations.** Estimated effect size maps showing the association between fluid (A,D) and crystallized (B,E) scores with relative CSA (A,B) and relative CTH (D,E). Maps are thresholded to show the largest 20% absolute beta estimates colored based on the sign of the association (red=positive; blue=negative). Similarity maps (C,F) show the overlap between the fluid and crystallized associations. Vertices with positive associations above threshold for both fluid and crystallized scores are shown in yellow and negative associations light blue (C,F).

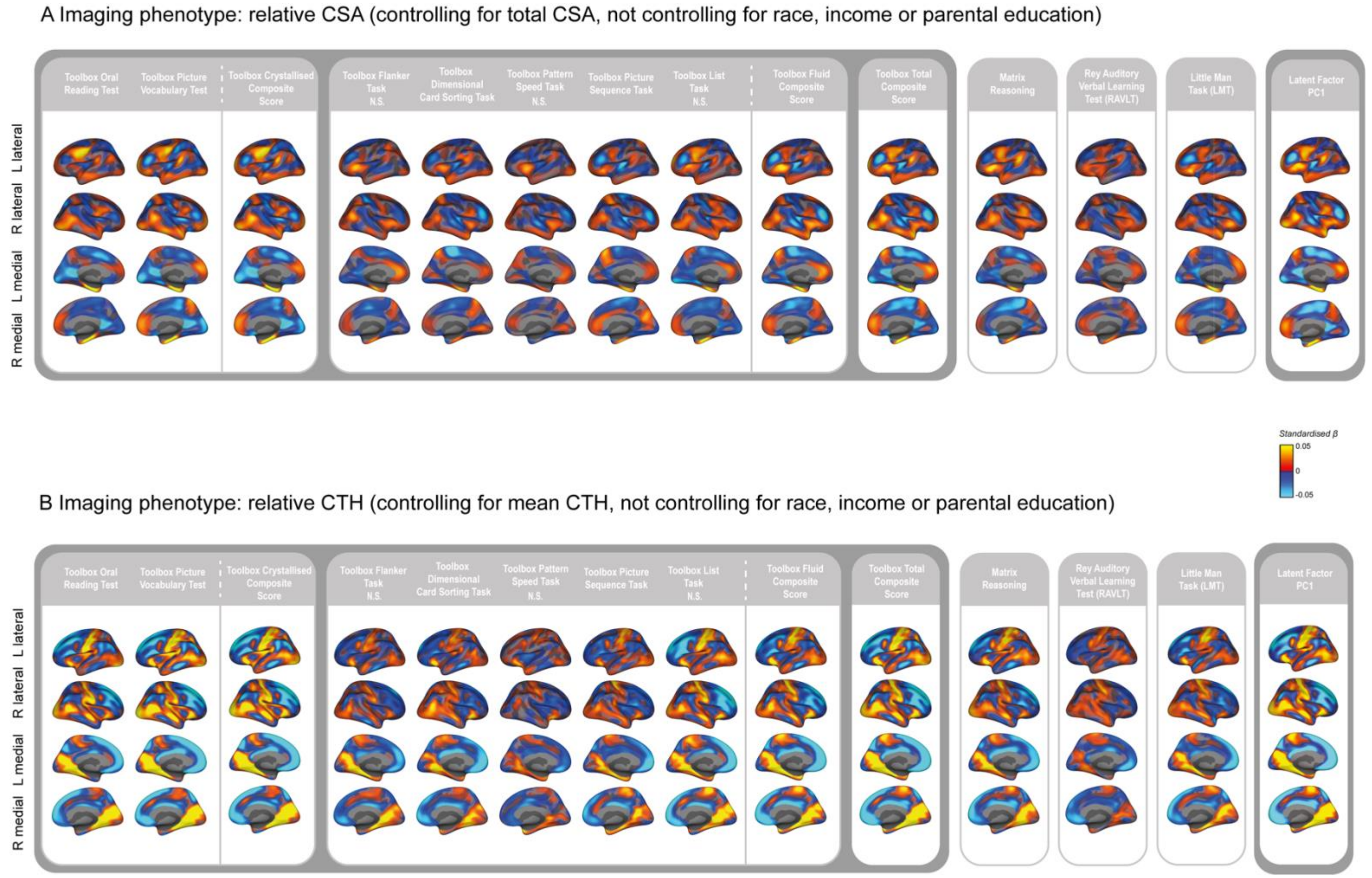


**Supplementary Figure 5.** Estimated effect size maps showing the mass univariate standardized beta coefficients for the association between each cognitive task and (A) the regionalization of CSA and (B) the regionalization of CTH without controlling for the sociodemographic variables of race/ethnicity, household income and parental education. In general, beta estimates were larger compared to when controlling for these variables (figure 4), particularly for relative CTH, and the effects were more similar across cognitive task.
